# L-Shaped Distributions of the Relative Substitution Rates (c/µ) in SARS-COV-2 with or without Molecular Clocks, Challenging Mainstream Evolutionary Theories

**DOI:** 10.1101/2024.04.29.591599

**Authors:** Chun Wu, Nicholas J. Paradis

**Affiliations:** Department of Chemistry and Biochemistry, Rowan University, Glassboro, NJ 08028, USA; Department of Biological & Biomedical Sciences, Rowan University, Glassboro, NJ 08028, USA

## Abstract

A definitive test to quantify fitness changes of mutations is required to end a continuing 50-year “neutralist-selectionist” debate in evolutionary biology. Our previous work introduced a substitution-mutation rate ratio c/µ test (c: substitution rate in Translated Region/TR or UnTranslated Region/UTR; µ: mutation rate) to quantify the selection pressure and thus the proportions of strictly neutral, nearly neutral, beneficial, and deleterious mutations in a genome. Intriguingly, both a L-shaped probability distribution of c/µ and molecular clock were observed for SARS-COV-2’s genome. We found that the proportion of the different mutation types from the distribution is not consistent with the hypotheses of the three existing evolution theories (Kimura’s Neutral Theory/KNT, Ohta’s Nearly Neutral Theory/ONNT and the Selectionist Theory/ST), and a balance condition explains the molecular clock, thus we proposed a new theory named as Near-Neutral Balanced Selectionist Theory (NNBST). In this study, the c/µ analysis was extended beyond the genome to 26 TRs, 12 UTRs, and 10 TRSs (Transcriptional Regulatory Sequences) of SARS-COV-2. While L-shaped probability distributions of c/µ were observed for all of 49 segments, molecular clocks were observed for only 24 segments, supporting NNBST and Near-Neutral Unbalanced Selectionist Theory (NNUST) to explain the molecular evolution of 24/25 segments with/without molecular clocks. Thus, the Near-Neutral Selectionist Theory (NNST) integrates traditional neutral and selectionist theories to deepen our understanding of how mutation, selection, and genetic drift influence genomic evolution.

**Author Summary:** The “neutralist-selectionist” debate in molecular evolution has been unresolved for 50 years due to the three main theories of molecular evolution (Kimura’s Neutral Theory/KNT, Ohta’s Nearly-Neutral Theory/ONNT, Selectionist Theory/ST) disagreeing on the proportion of neutral mutations (KNT), nearly-neutral deleterious mutations (ONNT) and adaptive mutations (ST) within species. We recently developed a robust method, the c/µ relative substitution rate test, to quantify the proportion of each mutation type within >11K genomic sequences of SARS-COV-2 RNA virus. Our previous analysis revealed an L-shaped c/µ probability distribution and a constant substitution rate (e.g., molecular clock) for the SARS-COV-2 genome over 19 months, and the proportions of mutation types were inconsistent with those predicted by the three theories. We thus proposed the Near-Neutral Balanced Selectionist Theory (NNBST) to explain the molecular clock-feature and L-shaped probability distribution for SARS-COV-2. In this study, we extended this analysis to the 25 protein-coding gene segments and 24 non-protein-coding segments of SARS-COV-2. We observed that all 49 segments exhibited an L-shaped probability distribution and 24 out of the 49 segments exhibited a molecular clock, however the remaining 25 segments did not exhibit a molecular clock. We thus propose the Near-Neutral Unbalanced Selectionist Theory (NNUST) and NNBST to explain the segments without/with molecular clock features, respectively. We also coin the Near-Neutral Selectionist Theory (NNST) to combine traditional KNT, ONNT and ST to deepen our understanding of how mutation, selection, and genetic drift influence genomic evolution.

## Introduction

All theories of molecular evolution (**Fig. 1A-B**) agree that most mutations are deleterious 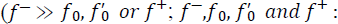: the fraction of deleterious, strictly neutral, nearly neutral and beneficial mutations) and thus will be purified through evolution, but they all differ in their hypotheses on the proportion of the strictly neutral, nearly neutral and beneficial mutations that become fixed in a population (A.K.A. substitution).

**Figure 1.**
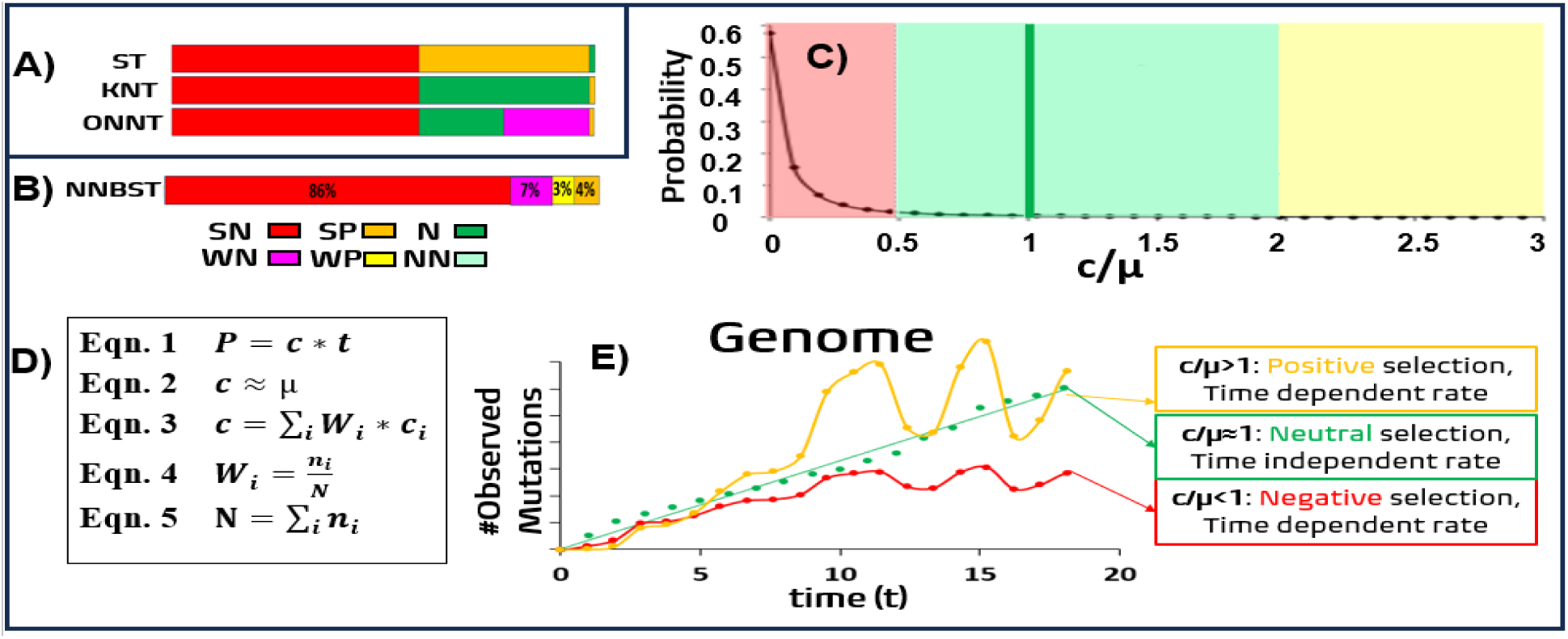
Comparison between the three mainstream evolution theories (A) and the newly proposed NNBST (B-E). **A:** Putative composition of the observed substitutions under different selection types for the three mainstream theories (ST, KNT and ONNT). **B:** The composition of SARS-COV-2 for NNBST. **SN:** strong negative selection (0 < c/µ < 0.5, red), **WN:** weak negative selection (0.5 < c/µ < 1.0, light red), **WP:** weak positive selection (1.0 < c/µ < 2.0, light yellow) and **SP:** strong positive selection (c/µ > 2.0, dark yellow). **NN**: WN + WP **C**: the L-shaped distribution of the relative substitution rate (c/µ) of SARS-COV-2. **D**: The five equations describing NNBST. **Equ. 1**: the integral substitution probability (**P**) per Nucleotide (NT) site of a genome at time *t* is the product of the substitution rate of the whole genome (c) with the time. **Equ. 2**: the substitution rate (c) is almost equal to the spontaneous mutation rate (µ) due to effective neutrality. **Equ. 3**: the substitution rate of the whole genome (c) is the weighted sum of the substitution rate of each segment (*c_i_*) with weight (*W_i_*). **Equ. 4**: The weight (*W_i_*) of each segment is determined by segment length (n_i_) over the genomic length (N). **Equ. 5**: the length of a genome (N) is a summation of the length of all segments. **E**: effective neutrality across the genome despite varying selective pressures acting on different segments.

The Selectionist (Neo-Darwinian) Theory/ST[1, 2]states that most substitutions are beneficial mutations that lead to a fitness increase of the species (i.e. *f*^+^ ≫ *f*_0_ ≈ 0). In contrast, Kimura’s Neutral Theory (KNT) [3, 4], [5, 6] argues that most substitutions are strictly neutral mutations that lead to no change in fitness (i.e. *f*_0_ ≫ *f*^+^ ≈ 0). Ohta’s Nearly Neutral Theory/ONNT[7, 8] extends this perspective by acknowledging the importance of nearly neutral mutations that have slightly (deleterious) effects 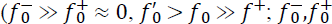: the fractions of slightly deleterious and slightly beneficial mutations) besides the strictly neutral mutations. Depending on the effective population size, environmental conditions, and selective pressures, these nearly neutral mutations can be fixed in the population with variable rates, likely leading to a time-dependent substitution rate. However, these theories could not be evaluated on the basis of empirical data if mutations could not be classified by their fitness changes into the different types and thus their proportion could not be quantified, thus leading to a continuing 50-year “neutralist-selectionist” debate[9] [10–12].

In our previous paper[13], we framed a replication-selection model (**Figure 2**) that phenomenologically quantifies selection pressure on any site in a genome including Translated Region (TR) and UnTranslated Region (UTR) using sequence data.

**Figure 2.**
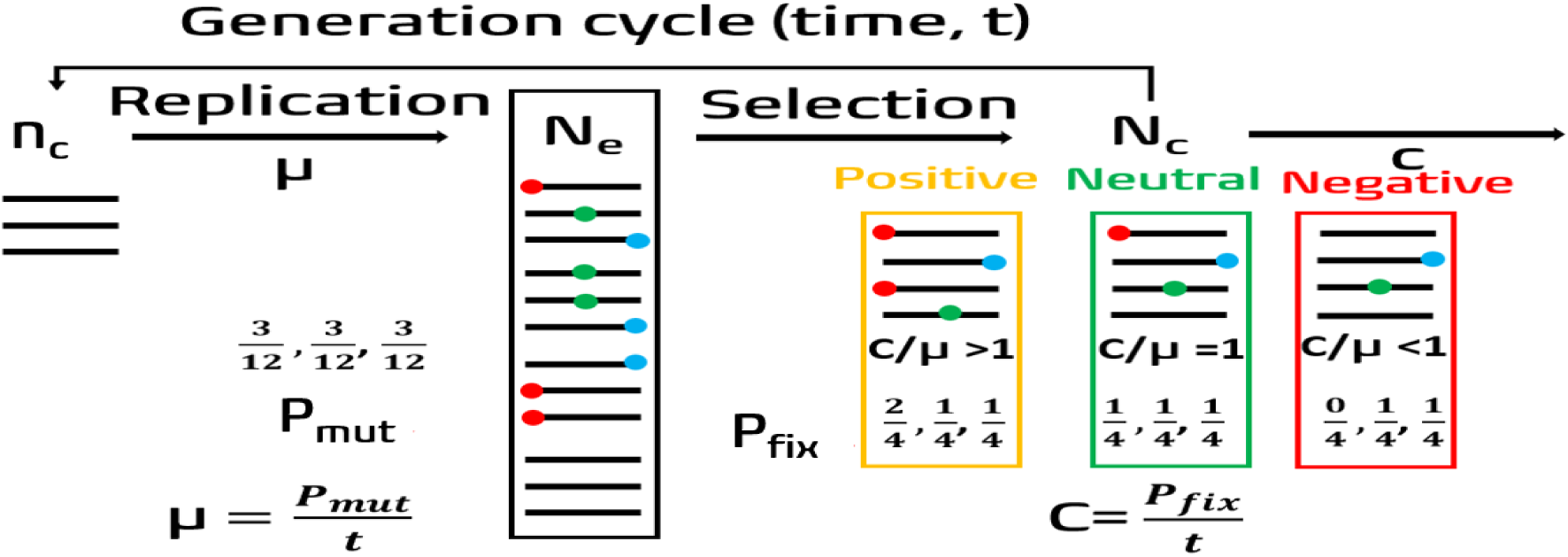
Simplified replication-selection model for a virus population over time under positive selection, neutral selection and negative selection for the first nucleotide position (red circle). A detailed description has been placed in the methods section. Viral genomes are shown as black lines. Mutations are shown as red, green and blue circles.

Our model uses a mutation-selection frame that assumes the mutation step is independent from the selection step[14]. Our model considers the substitution rate (c) in the population as a product of the mutation rate (µ) with a substitution-mutation rate ratio (c/µ), reflecting neutral and natural selection effects. Therefore, this c/µ ratio (c: substitution rate in TR or UTR, µ: mutation rate) can serve as a test to detect fitness change of mutations in a genome (c/µ = 1, <1 and >1: strictly neutral, deleterious, and beneficial). The relative substitution rate c/µ for each nucleotide (NT) site in the genome of SARS-COV-2 (29,903 NTs) was obtained from its first 19-month genomic data (three independent sets of a total of 11,198 genome sequences from NextStrain[15]) to quantify the fitness change of mutations for the very first time (**Fig. 1B-C**). Intriguingly, both L-shaped discrete probability distributions of c/µ (**Fig. 1C**) and molecular clock (**Fig. 1E**) were observed for the whole genome. From the distribution, we found that the proportion of strictly neutral, nearly neutral, beneficial mutations are not consistent with the hypotheses of the three existing theories. We therefore proposed a novel theory that merges some aspects of ONNT and ST to explain the observed mutation composition for the genome (86% strongly negative, 7% weakly negative, 3% weakly positive, 4% strongly positive). We named this new theory as Near-Neutral Balanced Selectionist Theory (NNBST). NNBST not only accounts for the important role of nearly neutral mutations (7% weakly negative and 3% weakly positive), but it also recognizes the crucial role from the positive selection (3% weakly positive and 4% strongly positive) in determining the overall substitution rate (**Fig. 1B-E**). NNBST challenges the core tenet of KNT and ONNT by suggesting that strictly neutral mutations are rare (0.5%, 1:20 between strictly neutral and near neutral mutations) due to natural selection, evidenced by the observation of an L-shaped distribution curve of c/µ rather than a Poisson distribution curve centered at strictly neutral (c/µ = 1) for SARS-COV-2 (**Fig. 1C**).

In addition, a balancing act between beneficial (c_i_/µ>1) and deleterious mutations (c_i_/µ<1) in increasing and decreasing the substitution rate respectively leads to a molecular evolutionary clock (i.e. an effective neutrality or c/µ = 1) across the pathogen’s genome despite varying selective pressures acting on different genes (**Fig. 1D-E**). This balanced natural selection mechanism leads to a molecular clock to describe the linear increase of genomic variation over time using a simple set of equations (**Fig. 1D**). This mechanism does not require the original stronger assumptions that the fraction of beneficial mutations are negligible by KNT to derive the molecular clock.[6] If the substitution rates between beneficial and deleterious mutations are not balanced, a molecular clock will not be observed. We coined it Near-Neutral Unbalanced Selectionist Theory (NNUST), which does not require the stronger assumption that that the fraction of near neutral mutations are negligible by ST. Combining NNBST with NNUST, a Near-Neutral Selectionist Theory (NNST) acknowledges the contribution of both near neutral and beneficial mutations in determining the overall substitution rate.[10]

In this study, the c/µ analysis was extended beyond the gnome to 26 TRs, 12 UTRs, and 10 TRSs (Transcriptional Regulatory Sequences) of SARS-COV-2. While discrete L-shaped probability distributions of c/µ were observed for all of 49 segments, molecular clocks (R^2^>0.6000) were observed for only 25 segments. Therefore, we propose NNBST and NNUST to explain the molecular evolution of 25 and 24 segments with and without showing molecular clocks, respectively. Our findings advocate for the c/µ test as an effective test, potentially ending the longstanding debate in evolutionary biology with NNST.

## Results

To validate c/µ values (per NT and per unit time) for 49 segments of SARS-COV-2 including 1 genome, 26 TRs, 12 UTRs and 10 TRSs, reported in our previous paper [13], that is based on position-based approach in which SRSRS (**Figure 4**) was used to get the rates (per 19 months per NT), their ratios, and the proportions of mutation types from cumulative PDRSR were calculated (**Figure 5**).

**Figure 3.**
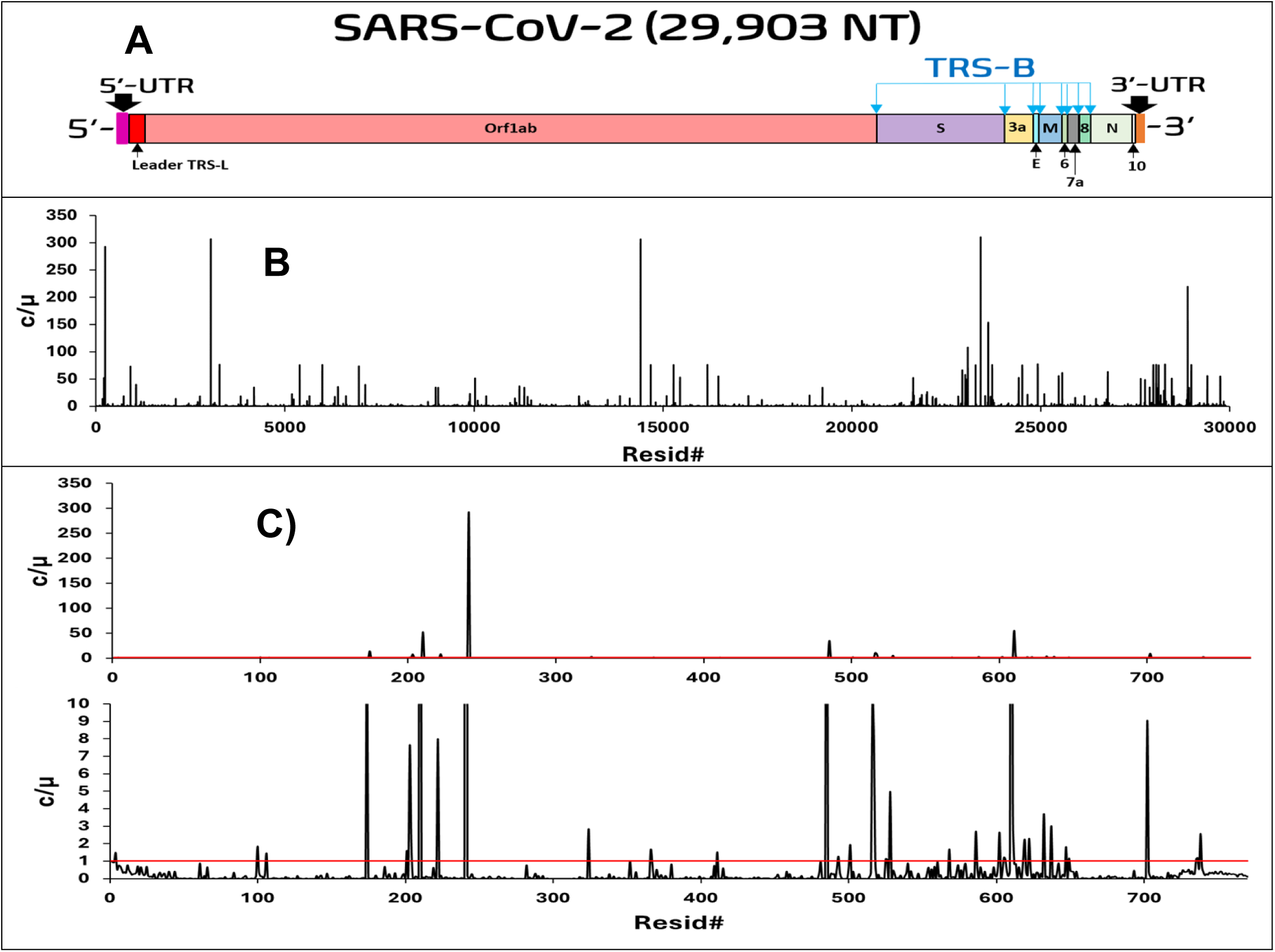
**A**) The structure of the SARS-CoV-2 genome, including major gene segments (Orf1ab, S, E, M and N), accessory genes (Orf3a, Orf6, Orf7a, Orf8 and Orf10), UTRs (Orf1ab 5’-UTR and Orf10 3’-UTR) and TRS (leader TRS-L and TRS-B). **B**) Position-based c/µ at each NT site in the SARS-COV-2 genome. **C**) Position-based c/µ at each NT site in All-UTR. The red line in Figure 3C represents neutral selection (c/µ=1). µ: the substitution rate of Orf1ab 5’UTR which is 5.46-fold of GSR.

**Figure 4.**
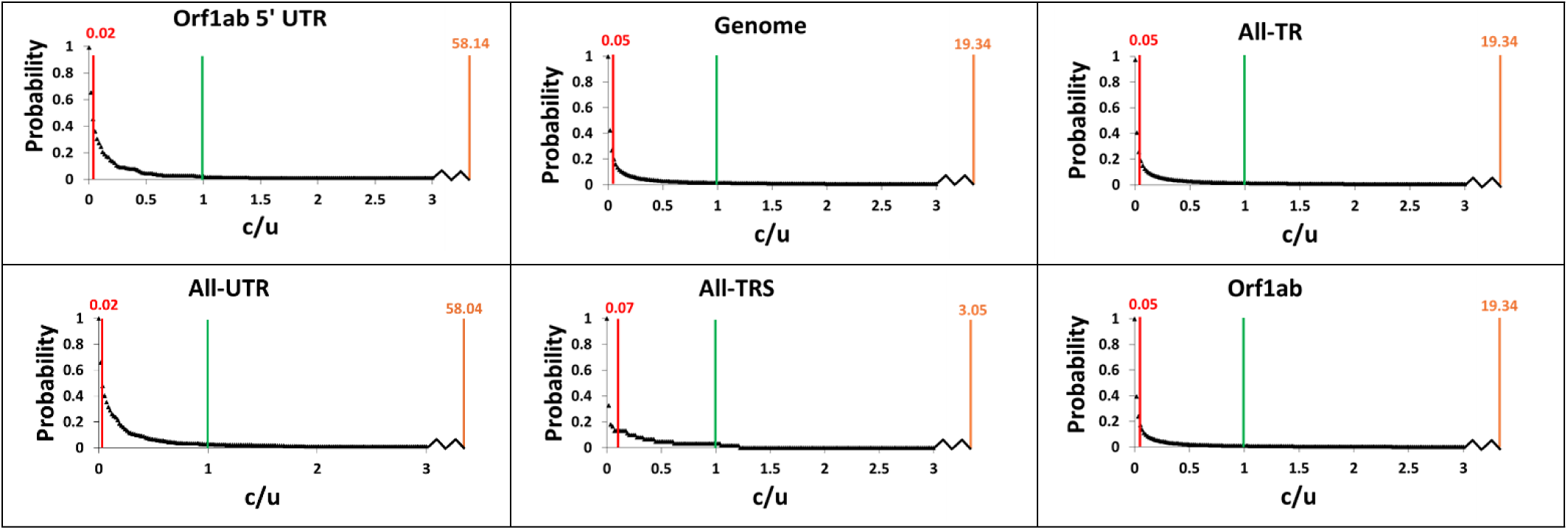
c/µ Cumulative probability distribution function of Orf1ab 5’UTR, genome, All-TR, All-UTR, All-TRS and Orf1ab segments demonstrating a high coefficient of determination, showing the abundance of sites under different selection types. The boundaries for weak negative (red line) to neutral selection (green line) to weak positive selection (orange line) and their determined c/µ positions are noted. Strong negative selection and strong positive selection would be to the left and right of the red and orange lines, respectively. A broken lined leading on the x-axis represents the distance between c/µ = 3.0 to the weak positive selection boundary. See **Figure S1** for the remaining segments demonstrating a high coefficient of determination.

These rates and ratios are also obtained in this study from the time-based approach in which temporal trend (timeline) in **Figure 5** is fitted by least square (y=m*x) to obtain the rates and the coefficient of determination (R^2^), thus their ratios from the three independent genome sequence datasets used in our previous paper as described in the method section (**Table 1** and **Figure S1-S3**). R^2^ offers a statistical measure on how good molecular clock feature (i.e. time-independent constant rate) holds.

**Figure 5.**
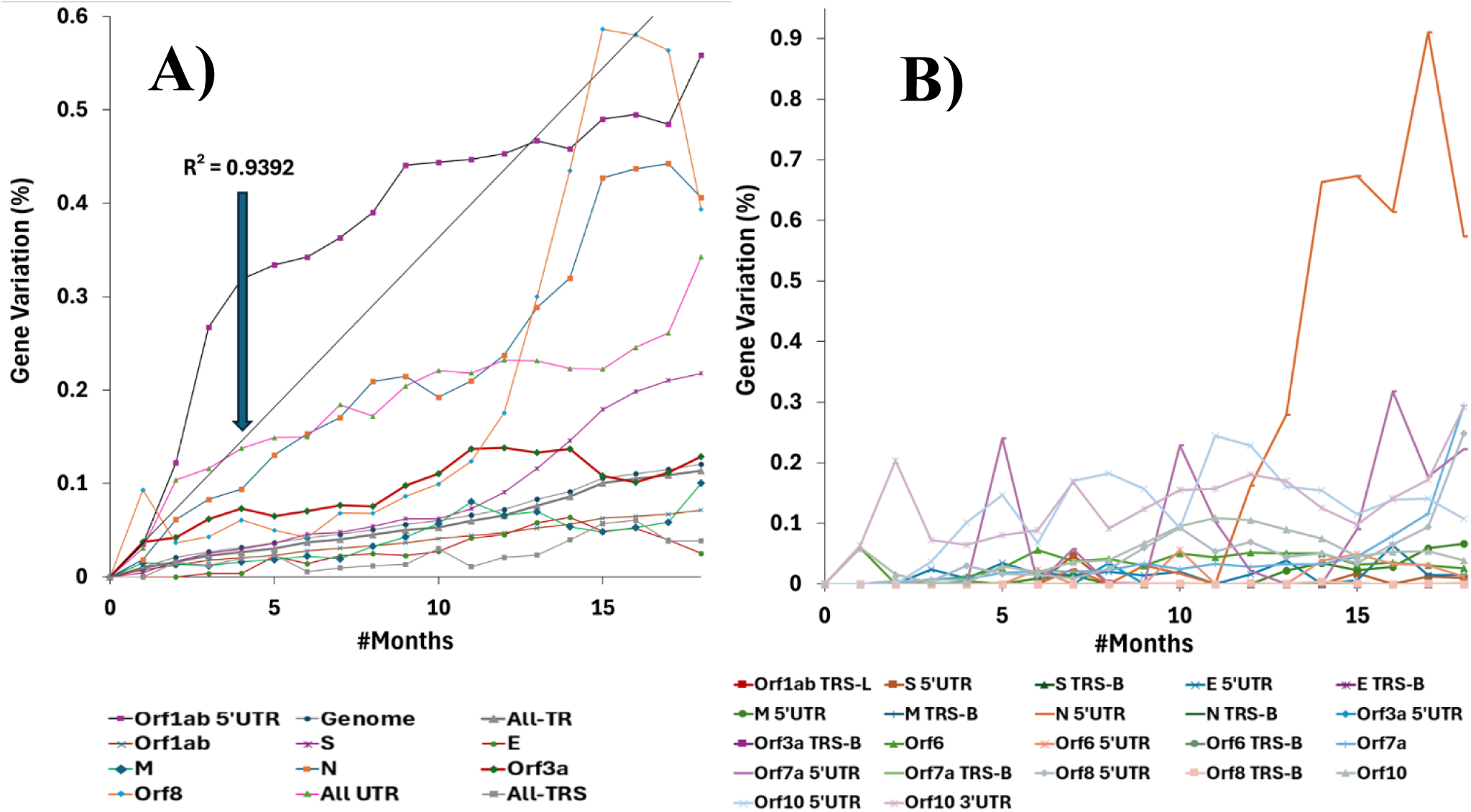
The total percent NT substitution variation (c) for segments exhibiting molecular clock (A) and non-molecular clock (B) features over evolution time, averaged over datasets A1A-A1C. The trendline and coefficient of determination for Orf1ab 5’UTR (R^2^ = 0.9392) is defined as µ. See **Figures S2-S3** for the individual timelines for each segment.

**Table 1.**
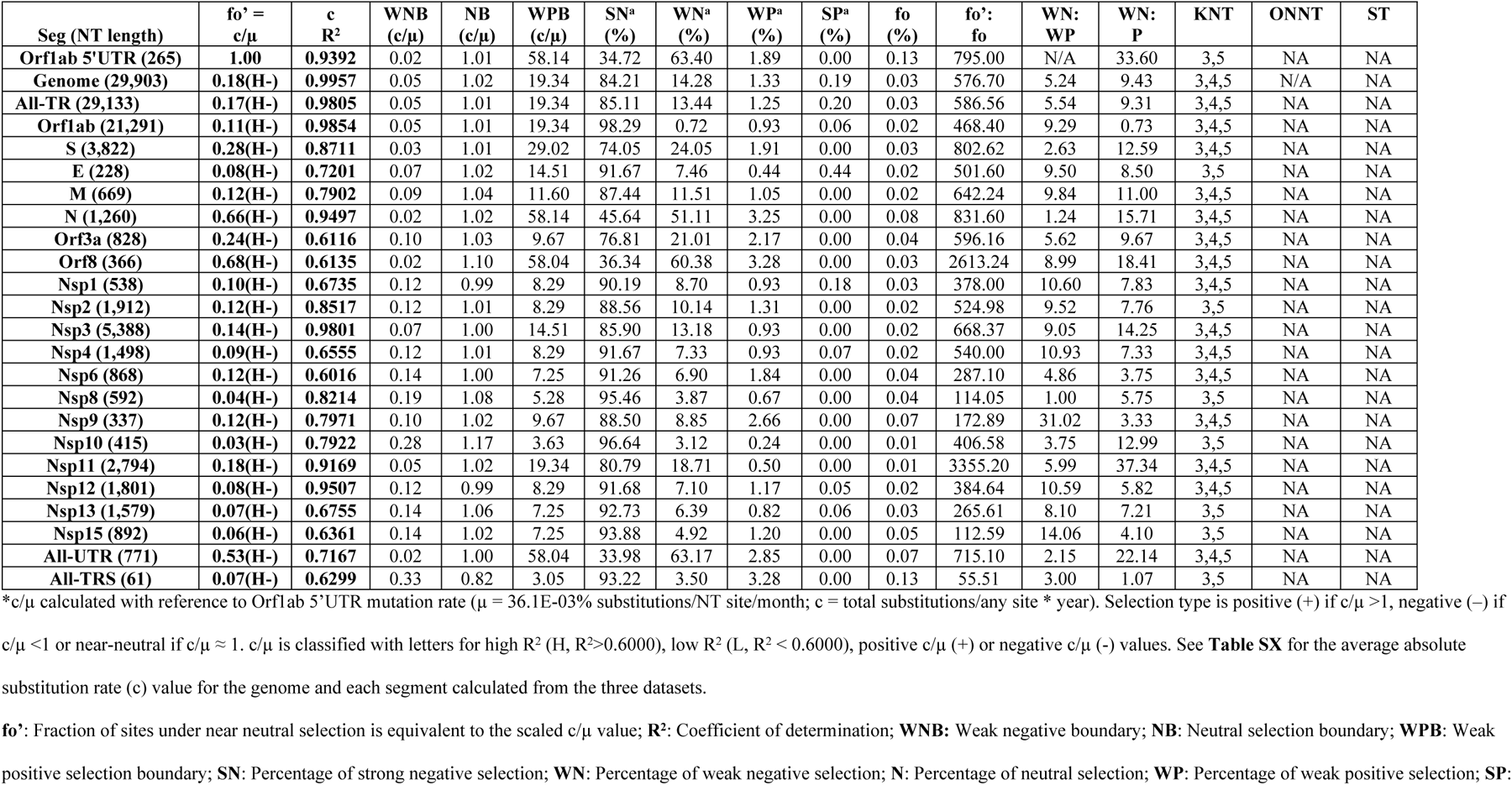
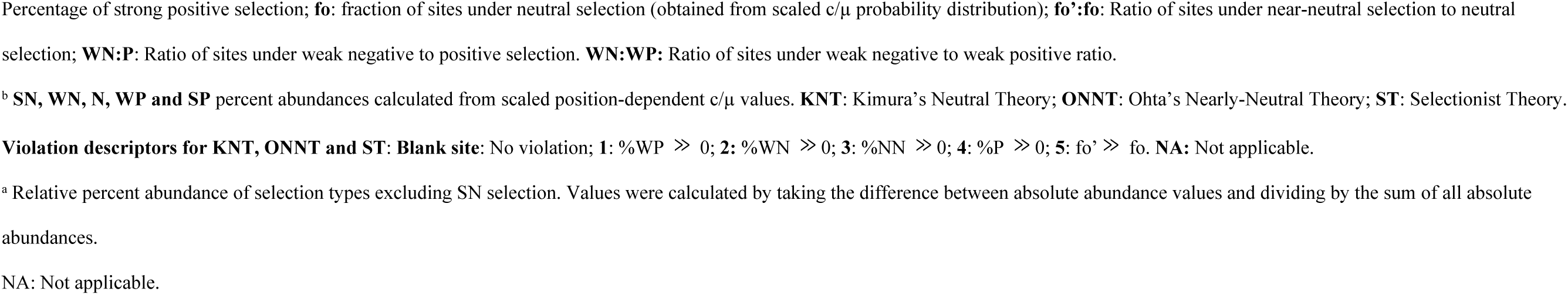
Time-based c/μ, R^2^, c/μ and percent abundance at three selection type boundaries, the relative percent abundance of sites under four different selection types, the ratio of sites under weak negative to positive selection, the ratio of sites under near-neutral to neutral selection and evaluation of KNT, ONNT and ST for the genome and each segment exhibiting molecular clock features of SARS-COV-2 over 19 months.

There are several earliest genome sequences such as Wuhan-Hu-1 and Wuhan IPBCAMS-WH-01 that can serve as the reference sequence for our substitution analysis. To quickly check if the substitution rate of the genome and the molecular clock feature change much when different reference sequences are used, we generated timelines using either as the reference sequence. The absolute rate changes slightly from 6.4E-3 to 7.2E-3 per NT per month and R^2^ is changed from 0.9829 to 0.9912 when the reference sequence is changed from Wuhan-Hu-1 to Wuhan IPBCAMS-WH-01 (**Figure S1**). Therefore, its effect on the relative substitution rate should be even smaller, and thus the former sequence which was used in our previous study was still used as reference sequence in this study.

### Molecular clock with R^2^>0.6000 was observed for only 24 out of 49 segments including the genome, UTR, TR, 5’UTR, and most of genes and NSPs of SARS-COV-2

The raw time-based c and c R^2^ values were tabulated (UTR/TRS: **Table S1; and** TR: **Table S2**). While the substitution timelines of Orf1ab 5’UTR, genome, All-TR, All-UTR, All-TRS and most of genes and NSPs of SARS-CoV-2 (24 segments in **Table 1**) have high coefficient of determination (R^2^>0.6000), most of UTR/TRS segments, a few of genes and NSPs (25 segments in **Table 2**) have low coefficient of determination (R^2^<0.6000).

**Table 2.**
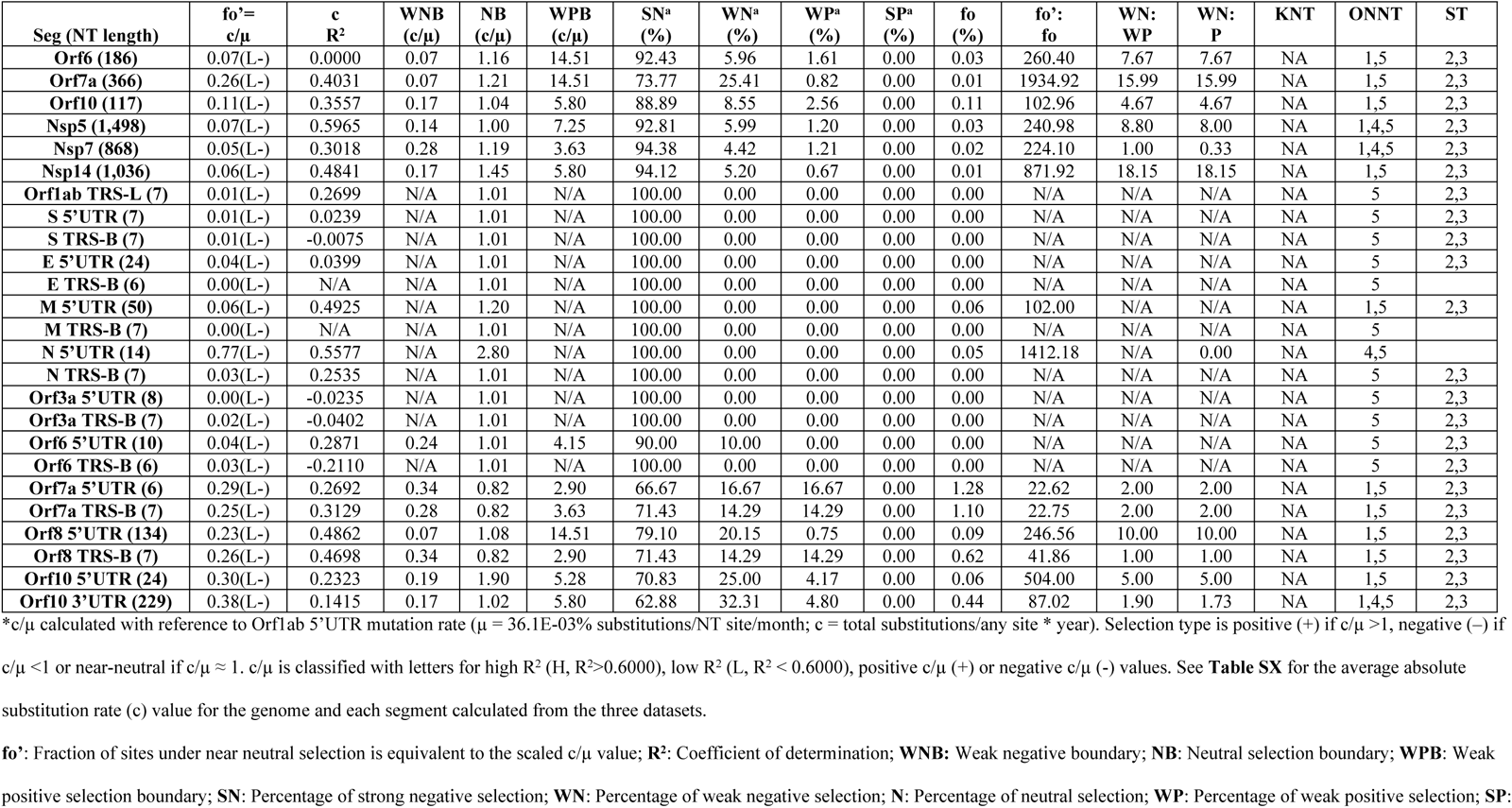
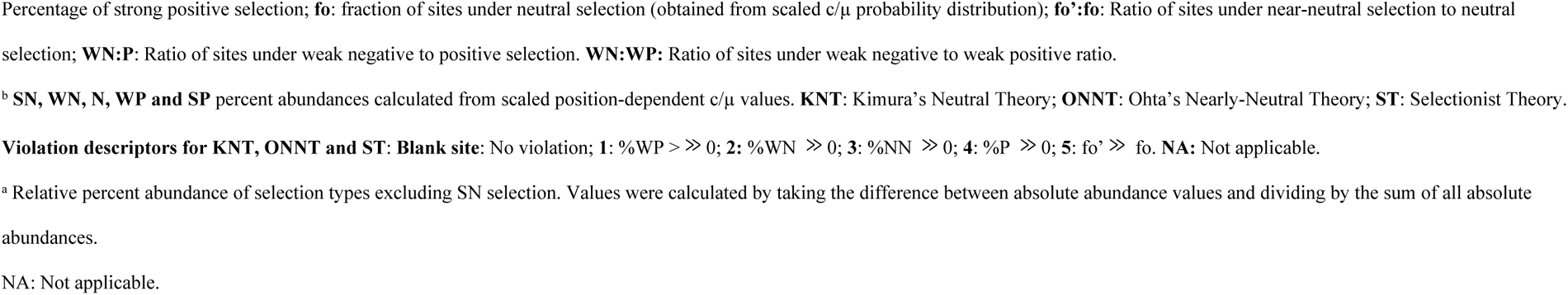
Position-based c/μ, R^2^, c/μ and percent abundance at three selection type boundaries, the relative percent abundance of sites under four different selection types, the ratio of sites under weak negative to positive selection, the ratio of sites under near-neutral to neutral selection and evaluation of KNT, ONNT and ST for each segment exhibiting non-molecular clock features of SARS-COV-2 over 19 months.

| Seg (NT length) | fo'=<br>c/$\mu$ | c<br>R<sup>2</sup> | WNB<br>(c/$\mu$) | NB<br>(c/$\mu$) | WPB<br>(c/$\mu$) | SN<sup>a</sup><br>(%) | WN<sup>a</sup><br>(%) | WP<sup>a</sup><br>(%) | SP<sup>a</sup><br>(%) | fo<br>(%) | fo':<br>fo | WN:<br>WP | WN:<br>P | KNT | ONNT | ST |
| --- | --- | --- | --- | --- | --- | --- | --- | --- | --- | --- | --- | --- | --- | --- | --- | --- |
| **Orf6 (186)** | 0.07(L-) | 0.0000 | 0.07 | 1.16 | 14.51 | 92.43 | 5.96 | 1.61 | 0.00 | 0.03 | 260.40 | 7.67 | 7.67 | NA | 1,5 | 2,3 |
| **Orf7a (366)** | 0.26(L-) | 0.4031 | 0.07 | 1.21 | 14.51 | 73.77 | 25.41 | 0.82 | 0.00 | 0.01 | 1934.92 | 15.99 | 15.99 | NA | 1,5 | 2,3 |
| **Orf10 (117)** | 0.11(L-) | 0.3557 | 0.17 | 1.04 | 5.80 | 88.89 | 8.55 | 2.56 | 0.00 | 0.11 | 102.96 | 4.67 | 4.67 | NA | 1,5 | 2,3 |
| **Nsp5 (1,498)** | 0.07(L-) | 0.5965 | 0.14 | 1.00 | 7.25 | 92.81 | 5.99 | 1.20 | 0.00 | 0.03 | 240.98 | 8.80 | 8.00 | NA | 1,4,5 | 2,3 |
| **Nsp7 (868)** | 0.05(L-) | 0.3018 | 0.28 | 1.19 | 3.63 | 94.38 | 4.42 | 1.21 | 0.00 | 0.02 | 224.10 | 1.00 | 0.33 | NA | 1,4,5 | 2,3 |
| **Nsp14 (1,036)** | 0.06(L-) | 0.4841 | 0.17 | 1.45 | 5.80 | 94.12 | 5.20 | 0.67 | 0.00 | 0.01 | 871.92 | 18.15 | 18.15 | NA | 1,5 | 2,3 |
| **Orf1ab TRS-L (7)** | 0.01(L-) | 0.2699 | N/A | 1.01 | N/A | 100.00 | 0.00 | 0.00 | 0.00 | 0.00 | N/A | N/A | N/A | NA | 5 | 2,3 |
| **S 5'UTR (7)** | 0.01(L-) | 0.0239 | N/A | 1.01 | N/A | 100.00 | 0.00 | 0.00 | 0.00 | 0.00 | N/A | N/A | N/A | NA | 5 | 2,3 |
| **S TRS-B (7)** | 0.01(L-) | -0.0075 | N/A | 1.01 | N/A | 100.00 | 0.00 | 0.00 | 0.00 | 0.00 | N/A | N/A | N/A | NA | 5 | 2,3 |
| **E 5'UTR (24)** | 0.04(L-) | 0.0399 | N/A | 1.01 | N/A | 100.00 | 0.00 | 0.00 | 0.00 | 0.00 | N/A | N/A | N/A | NA | 5 | 2,3 |
| **E TRS-B (6)** | 0.00(L-) | N/A | N/A | 1.01 | N/A | 100.00 | 0.00 | 0.00 | 0.00 | 0.00 | N/A | N/A | N/A | NA | 5 |  |
| **M 5'UTR (50)** | 0.06(L-) | 0.4925 | N/A | 1.20 | N/A | 100.00 | 0.00 | 0.00 | 0.00 | 0.06 | 102.00 | N/A | N/A | NA | 1,5 | 2,3 |
| **M TRS-B (7)** | 0.00(L-) | N/A | N/A | 1.01 | N/A | 100.00 | 0.00 | 0.00 | 0.00 | 0.00 | N/A | N/A | N/A | NA | 5 |  |
| **N 5'UTR (14)** | 0.77(L-) | 0.5577 | N/A | 2.80 | N/A | 100.00 | 0.00 | 0.00 | 0.00 | 0.05 | 1412.18 | N/A | 0.00 | NA | 4,5 |  |
| **N TRS-B (7)** | 0.03(L-) | 0.2535 | N/A | 1.01 | N/A | 100.00 | 0.00 | 0.00 | 0.00 | 0.00 | N/A | N/A | N/A | NA | 5 | 2,3 |
| **Orf3a 5'UTR (8)** | 0.00(L-) | -0.0235 | N/A | 1.01 | N/A | 100.00 | 0.00 | 0.00 | 0.00 | 0.00 | N/A | N/A | N/A | NA | 5 | 2,3 |
| **Orf3a TRS-B (7)** | 0.02(L-) | -0.0402 | N/A | 1.01 | N/A | 100.00 | 0.00 | 0.00 | 0.00 | 0.00 | N/A | N/A | N/A | NA | 5 | 2,3 |
| **Orf6 5'UTR (10)** | 0.04(L-) | 0.2871 | 0.24 | 1.01 | 4.15 | 90.00 | 10.00 | 0.00 | 0.00 | 0.00 | N/A | N/A | N/A | NA | 5 | 2,3 |
| **Orf6 TRS-B (6)** | 0.03(L-) | -0.2110 | N/A | 1.01 | N/A | 100.00 | 0.00 | 0.00 | 0.00 | 0.00 | N/A | N/A | N/A | NA | 5 | 2,3 |
| **Orf7a 5'UTR (6)** | 0.29(L-) | 0.2692 | 0.34 | 0.82 | 2.90 | 66.67 | 16.67 | 16.67 | 0.00 | 1.28 | 22.62 | 2.00 | 2.00 | NA | 1,5 | 2,3 |
| **Orf7a TRS-B (7)** | 0.25(L-) | 0.3129 | 0.28 | 0.82 | 3.63 | 71.43 | 14.29 | 14.29 | 0.00 | 1.10 | 22.75 | 2.00 | 2.00 | NA | 1,5 | 2,3 |
| **Orf8 5'UTR (134)** | 0.23(L-) | 0.4862 | 0.07 | 1.08 | 14.51 | 79.10 | 20.15 | 0.75 | 0.00 | 0.09 | 246.56 | 10.00 | 10.00 | NA | 1,5 | 2,3 |
| **Orf8 TRS-B (7)** | 0.26(L-) | 0.4698 | 0.34 | 0.82 | 2.90 | 71.43 | 14.29 | 14.29 | 0.00 | 0.62 | 41.86 | 1.00 | 1.00 | NA | 1,5 | 2,3 |
| **Orf10 5'UTR (24)** | 0.30(L-) | 0.2323 | 0.19 | 1.90 | 5.28 | 70.83 | 25.00 | 4.17 | 0.00 | 0.06 | 504.00 | 5.00 | 5.00 | NA | 1,5 | 2,3 |
| **Orf10 3'UTR (229)** | 0.38(L-) | 0.1415 | 0.17 | 1.02 | 5.80 | 62.88 | 32.31 | 4.80 | 0.00 | 0.44 | 87.02 | 1.90 | 1.73 | NA | 1,4,5 | 2,3 |

\* $c/\mu$ calculated with reference to Orf1ab 5'UTR mutation rate ( $\mu = 36.1E-03\%$ substitutions/NT site/month; $c$ = total substitutions/any site \* year). Selection type is positive (+) if $c/\mu > 1$ , negative (-) if
$c/\mu < 1$ or near-neutral if $c/\mu \approx 1$ . $c/\mu$ is classified with letters for high $R^2$ (H, $R^2 > 0.6000$ ), low $R^2$ (L, $R^2 < 0.6000$ ), positive $c/\mu$ (+) or negative $c/\mu$ (-) values. See **Table SX** for the average absolute substitution rate ( $c$ ) value for the genome and each segment calculated from the three datasets.
**fo'**: Fraction of sites under near neutral selection is equivalent to the scaled $c/\mu$ value; **R<sup>2</sup>**: Coefficient of determination; **WNB**: Weak negative boundary; **NB**: Neutral selection boundary; **WPB**: Weak positive selection boundary; **SN**: Percentage of strong negative selection; **WN**: Percentage of weak negative selection; **N**: Percentage of neutral selection; **WP**: Percentage of weak positive selection; **SP**:
Percentage of strong positive selection; **fo**: fraction of sites under neutral selection (obtained from scaled $c/\mu$ probability distribution); **fo':fo**: Ratio of sites under near-neutral selection to neutral selection; **WN:P**: Ratio of sites under weak negative to positive selection. **WN:WP**: Ratio of sites under weak negative to weak positive ratio.
<sup>b</sup> **SN, WN, N, WP and SP** percent abundances calculated from scaled position-dependent $c/\mu$ values. **KNT**: Kimura's Neutral Theory; **ONNT**: Ohta's Nearly-Neutral Theory; **ST**: Selectionist Theory.
**Violation descriptors for KNT, ONNT and ST: Blank site**: No violation; **1**: %WP > >> 0; **2**: %WN >> 0; **3**: %NN >> 0; **4**: %P >> 0; **5**: fo' >> fo. **NA**: Not applicable.
<sup>a</sup> Relative percent abundance of selection types excluding SN selection. Values were calculated by taking the difference between absolute abundance values and dividing by the sum of all absolute abundances.
NA: Not applicable.

For the UTR, c R^2^ values revealed only 3 out of 22 non-coding segments exhibited molecular clocks. Orf1ab 5’UTR (0.9392), All-UTR (0.7167) and All-TRS (0.6299) exhibited good molecular clocks, whereas the remaining UTRs and TRSs did not exhibit good molecular clocks (-0.2110 - 0.5577), with E TRS-B and M TRS-B being entirely conserved (c/μ=0) and thus no R^2^ was generated (**Table S1).** For the TR, c R^2^ values demonstrated 20 out of 25 coding segments showed good molecular clocks. Besides the genome (0.9957) and All-TR (0.9805), Orf1ab (0.9854), N (0.9497), S (0.8711), M (0.7902), E (0.7201), Orf8 (0.6135), Orf3a (0.6116), NSP1-4 (0.6555-0.9801), NSP6-13 (0.6755-0.9507) and NSP15 (0.6361) exhibit good molecular clocks, whereas NSP5 (0.5965), NSP14 (0.4841), Orf7a (0.4031), Orf10 (0.3557), NSP7 (0.3018) and Orf6 (0.0000) do not exhibit molecular clock features (**Table S2**).

### The highest substitution rate of Orf1ab 5’UTR with a molecular clock was used as the Fundamental Genomic Mutation Rate (*µ*) of SARS-CoV-2

In our previous study[13], µ was its genomic substitution rate (GSR) derived from **equation 8a** (*c* = μ*f*^′^_0_ ). GSR was obtained from least square fitting of a linear line of the genomic substitutions over the time with a high correlation coefficient (R^2^=0.9957) using **equation 1** (*P* = *c* ∗ *t*). This (μ = *c*) sets the fraction of near neutral mutation sites in the genome be one (*f*^′^_0_ = 1). However, the substitution rates of some segments (e.g. Orf1ab 5’UTR, Orf8, N, all-UTR, S and Orf3a in decreasing order) also show a time-independent molecular clock feature (R^2^>0.6000) with a much higher rate than GSR (e.g. 5.46-fold of GSR for Orf1ab 5’UTR). If GSR is still used as µ, the fraction of near neutral mutation sites in Orf1ab 5’UTR would go beyond one. To remove this kind of violation, the highest substitution rate of the segment satisfying molecular clock was set to be µ. In this study, the substitution rate of Orf1ab 5’UTR was used as µ.

### Time-based and position-based c/μ are highly consistent for the 24 segments with molecular clock feature, also suggesting validity of molecular clock for these segments

The consistency of time-based and position-based c/µ values were assessed for UTR/TRS segments (**Table S3**) and the TR segments (**Table S4)** to verify the accuracy of the molecular clock assumption in the time-based approach. The consistency of both methods is dependent on the linearity (R^2^) of c obtained from least square fitting: substitution rates exhibiting a good molecular clock (R^2^>0.6000) should give very consistent c/µ values, whereas the absence of a molecular clock will produce larger deviations in the time-based and position-based c/µ as descripted in our method section. The absolute and percent differences between time-based and position-based c/µ methods were assessed to check the robustness of c/µ obtained from linear regression. For the UTR/TRS, larger differences were observed in segments with good molecular clocks (0.03-0.18, or 1.74%-8.11%) and without molecular clocks (0.00-0.71, or 0.00%-55.56%). For the TR, smaller differences were observed for segments exhibiting molecular clocks (0.00-0.07, or 0.00%-12.50%) and larger differences were observed in segments without molecular clocks (0.01-0.54, or 0.53%-38.57%). In summary, despite the larger percent differences of some segments (e.g., Orf6, Orf7a, Orf10, Orf1ab TRS-L, Orf3a TRS-B, etc.), 21 out of 25 coding segments and only All-UTR, All-TRS and Orf1ab 5’UTR exhibited a good molecular clock, indicating their selection type assignment by time-based c/µ is consistent with the position-based c/µ values. The remaining four coding segments (Orf7a, Orf10, NSP5 and NSP7) and each UTR and TRS should use the position-based c/µ for their accurate selection type assignment.

### Discrete L-shaped probability distributions of c/µ were observed for all of 49 segments

Our previous paper [13] has shown that an L-shaped distribution curve rather than a Poisson distribution curve centered at 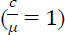 was observed for all 49 segments (**Figures S6-S8** [13]). The c/µ of these discrete distributions was scaled down by 5.46-fold to reflect the new µ in this study (**Figure S4-S5**). To facilitate the assessment for different mutation types, the cumulative probability distribution function 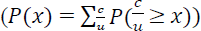 was also generated for all 49 segments (**Figure 5** and **Figure S6-S7**). Although the c/µ scaling does not change the fraction of conserved sites 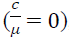, it does change the absolute fraction of strictly neutral, nearly neutral, beneficious, and deleterious mutation sites. The proportion of the strict neutral, nearly neutral, beneficial, deleterious mutations were determined from L-shaped PDRSR as described in the methods section (**Table 3-4 and Figure 7**).

**Figure 7.**
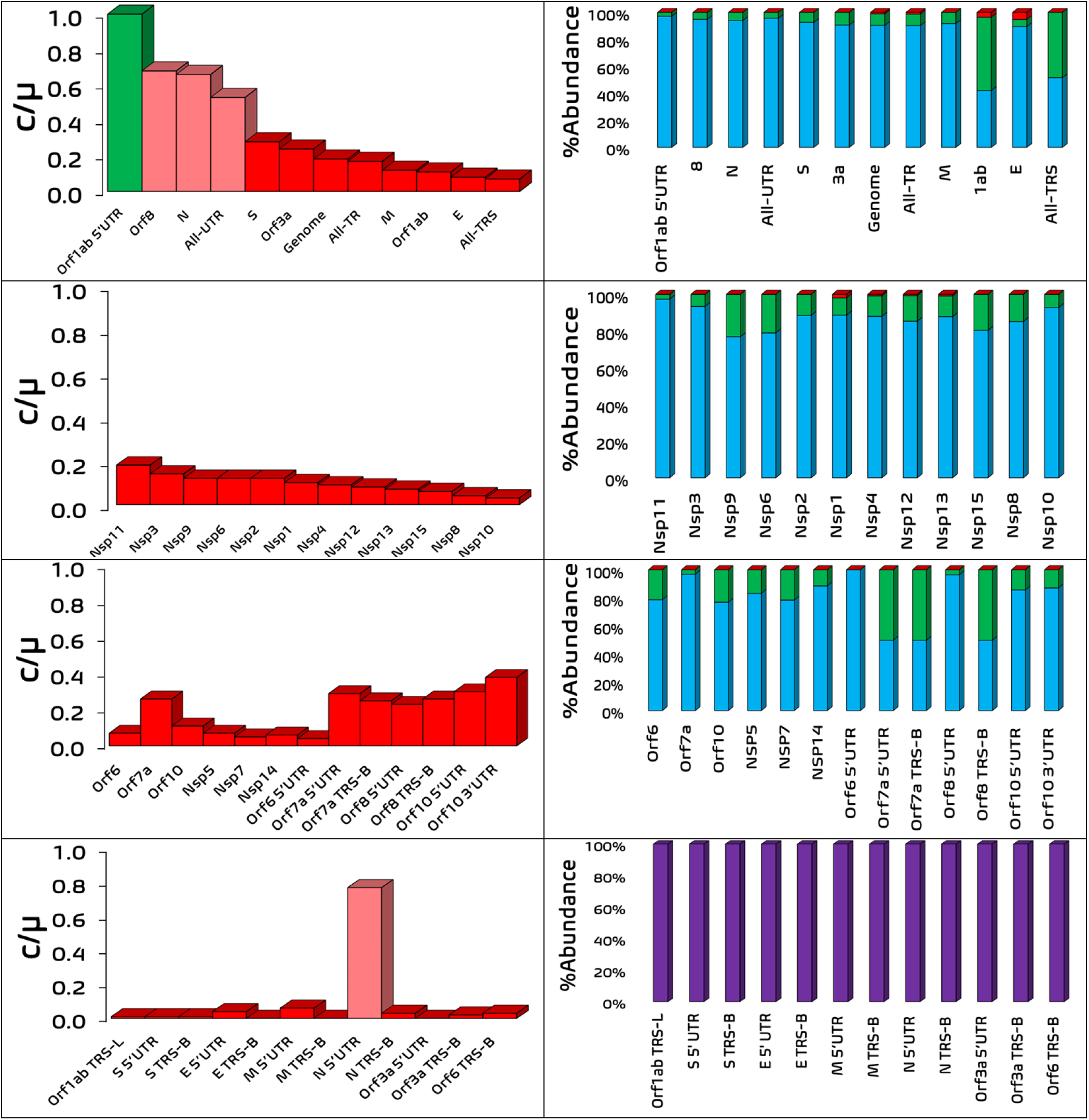
c/µ values (left column) and abundance of selection type (right column) for segments exhibiting molecular clock features (first and second rows) and non-molecular clock features (third and fourth rows). (Left column) Green, light red and dark red represent segments under neutral selection, weak negative selection and strong negative selection, respectively. (Right column) Blue, green, red and purple (where applicable) colors represent percentage of sites under weak negative, weak positive, strong positive and strong negative selection, respectively See **Figures S6-S7** for the graphs of segments decomposed into other selection types. See **Table S5** for tabulated values.

### Negligible fraction of strictly neutral mutation sites (*f*_0_ ≈ 0) suggests strictly neutral mutations due to genetic drifting are very rare and non-neutral mutations due to natural selection are predominant

The discrete distribution allows us to precisely determine the fraction of strictly neutral mutations 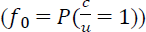. For the 24 segments with molecular clock, *f*_0_ ranges from 0.03% of the genome to 1.64% of All-TRS (**Table 3**). For the 25 segments without molecular clock, *f*_0_ ranges from 0.00% of 7 TRSs and 8 UTRs to 16.67% of Orf7a 5’UTR, 14.29% of Orf7a and Orf8 TRS-B, 0.85% of Orf10, 0.75% of Orf8 5’, 0.54% of Orf6, 0.44% of Orf10 3’UTR, 0.40% of Nsp7, and 0.05% of Nsp5 (**Table 4**). Therefore, most of the mutations are non-neutral mutations due to natural selection.

### The ratio of nearly neutral to strictly neutral mutations is greater than 2 up to 817 for 34 applicable segments, supporting that nearly neutral mutations play a more important role than strictly neutral mutations

This *f*^′^_0_:*f*_0_ ratio of the genome is ∼577. For the 24 segments with molecular clock, this ratio spans from ∼4 of All-TRS, to ∼468 of Orf1ab, to ∼587 of All-TR, and to ∼817 of All-UTR (**Table 3**). For the 25 segments with molecular clock, this ratio is only available for 10 segments, ranging from ∼2 of Orf7a 5’UTR, Orfa TRS-B and Orf8 TRS-B; to ∼13 of Orf6, Orf10, and Nsp7; to ∼31 of Orf8 5’UTR; to 87 of Orf10 3’UTR; to 129 of Nsp5; and 190 of Orf7a (**Table 4**).

### The ratio of mutations under weak negative to weak positive selection is greater than 1 up to 31 for 36 applicable segments, suggesting that although weak negative sites are more abundant than weak positive sites, weak positives do exist

This WP:WN ratio of the genome is ∼5. For the 24 segments with molecular clock, this ratio is available for 23 segments, spanning from ∼1 of Nsp8 and N; ∼2 of All-UTR; to ∼3 of S and All-TRS; to 5-6 of All-TR, Orf3a, Nsp6, and Nsp11; to 9-11 of Orf1ab, E, M, Orf8, Nsp2-4, and Nsp12; to ∼14 of Nsp15; and 31 of Nsp9 (**Table 3**). For the 25 segments with molecular clock, this ratio is only available for 12 segments, ranging from ∼1-2 of Orf10 3’UTR, Orf8 TRS-B, Orf7a TRS-B, Orf7a 5’UTR, and Nsp7; to ∼5-9 of Orf10 5’UTR, Orf6, Nsp5, Orf8 5’UTR and Orf10; to ∼16-18 of Orf7a and Nsp14 (**Table 4**).

### The ratio of mutations under weak negative to positive selection is greater than 1 up to 37 for all 37 applicable segments except for Orflab (0.73) and NSP7 (0.33) and Orf8 TRS-B (1.00), suggesting although more substitutions result from random fixation of slightly deleterious mutations, the beneficial substitutions due to positive selection are not so rare and could be more than slightly deleterious mutations for a few segments

This WN:P ratio of the genome is ∼9. For the 24 segments with molecular clock, its spans from 0.73 of Orflab to 37.34 of Nsp11 with a mean of 11±9 (**Table 3**). For the 25 segments with molecular clock, this ratio is only available for 12 segments, ranging from 0.33 of Nsp7 to 18.15 with a mean of 6.4±5.9 (**Table 4**). This ratio is almost identical to its WN:WP ratio, because there is almost no SP site for many of them.

### While NNST (union of NNBST and NNUST) are consistent with 6 empirical features of these 49 segments, KNT, ONNT and ST have many violations

The molecular clock feature (Yes or No), the abundance of weak positive sites (criteria 1: %WP ≫ 0), the abundance of weak negative sites (criteria 2: %WN ≫ 0), the abundance of nearly neutral sites (criteria 3: %NN ≫ 0), the abundance of positive sites (criteria 4: %P ≫ 0), and the ratio of nearly neutral to strictly neutral sites (criteria 5: fo’≫ fo) of 49 segments (**Table 3-4**) are used to evaluated the five evolutionary theories in **Table 3-4** (KNT, ONNT, ST, NNBST and NNUST). The summary of this evaluation is tabulated in **Table 5**.

**Table 5.**
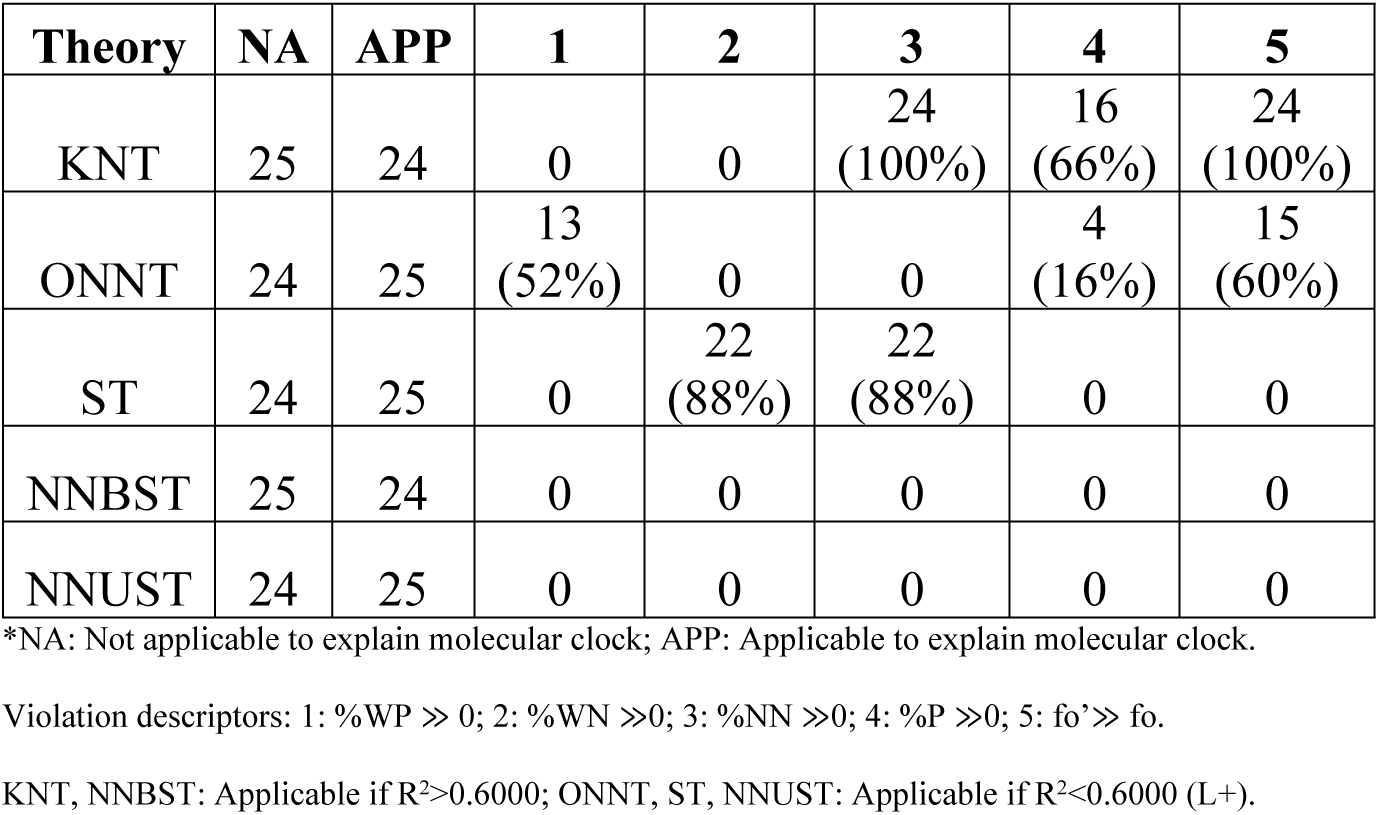
Evaluation of KNT, ONNT, ST, NNBST and NNUST for all segments. See **Table S9** for the evaluation at each segment. The number and percentage of violations calculated for applicable segments are presented.

KNT is not consistent with the lack of molecular clock for the 25 segments (**Table 4**). For the 24 segments with molecular clock, KNT is not consistent with criteria 3, 4 and 5 in 100%, 66%, and 100% cases, respectively (**Table 3&5**). In other words, KNT overestimates the abundance of strictly neutral sites (criteria 5), underestimates the abundance of near neutral sites (criteria 3) and positive sites (criteria 4). ONNT is not consistent with molecular clock for the 24 segments (**Table 3**). For the 25 segments without molecular clock, ONNT is not consistent with criteria 1, 4 and 5 in 52%, 16% and 60% cases, respectively (**Table 4 & 5**). In other words, ONNT overestimates the abundance of strictly neutral sites (criteria 5), underestimates the abundance of weak positive sites (criteria 1) and positive sites (criteria 4). ST is not consistent with molecular clock for the 24 segments (**Table 3**). For the 25 segments without molecular clock, ST is not consistent with criteria 2 and 3 in 88% and 88% cases, respectively (**Table 4 & 5**). In other words, ST underestimates the abundance of weak negative sites (criteria 2) and near neutral sites (criteria 3). While NNBST is not consistent with the lack of molecular clock for the 25 segments (**Table 4**), NNUST is not consistent with molecular clock for the 24 segments (**Table 3**). Without the strong assumptions in KNT, ONNT and ST, NNBST and NNUST have no violation on criteria 1-5.

### Scaling up of µ likely increases WN:P and *f*^′^_0_:*f*_0_ for the genome

What if the true µ is higher than the substitution rate of Orf1ab 5’UTR (5.46-fold of GSR)? Given the cumulative probability distribution of the genome, the change of WN:P and *f*^′^_0_:*f*_0_ can be determined when µ is scaled up beyond the 5.46 scaling factor (µ’). The scaled-up µ’ is determined from the cumulative probability of the original distribution with GSR as 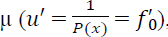, leading to determination of WNB and WPB; the factions of strictly neutral, near neutral, weak negative, weak positive; and the two ratios (**Figure 8**).

**Figure 8.**
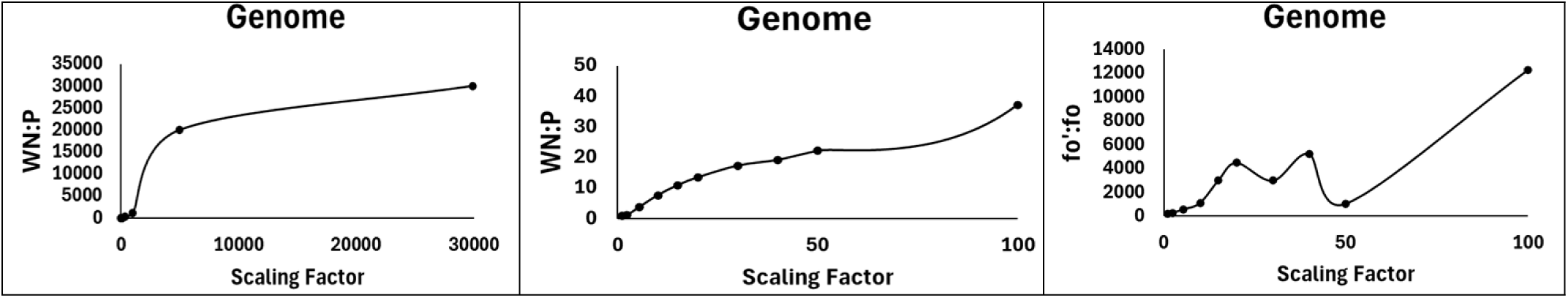
The impact of c/µ scaling factors on WN:P (left and middle) and fo’:fo (right) parameters. See **Tables S7-S8** for tabulated parameters.

As µ’ increases, the WN:P and *f*^′^_0_:*f*_0_ ratios of the gnome increase monotonically, being more consistent with NNBST than KNT and ST.

### The balancing condition explains molecular clock feature of a polyprotein gene (Orf1ab) of SARS-CoV-2

The balance between the sites of higher substitution rate under more positive selection and the sites of lower rate under more negative selection explained the observed molecular clock at the genome level of SARS-CoV-2 in our previous study[13]. This mechanism also works at a segment level for its polyprotein gene (Orf1ab) including 15 NSPs. The monthly total NT percent variation of NSP-15 and Orf1ab was plotted together to show which NSP is under more positive or negative selection (**Figure 9**).

**Figure 9.**
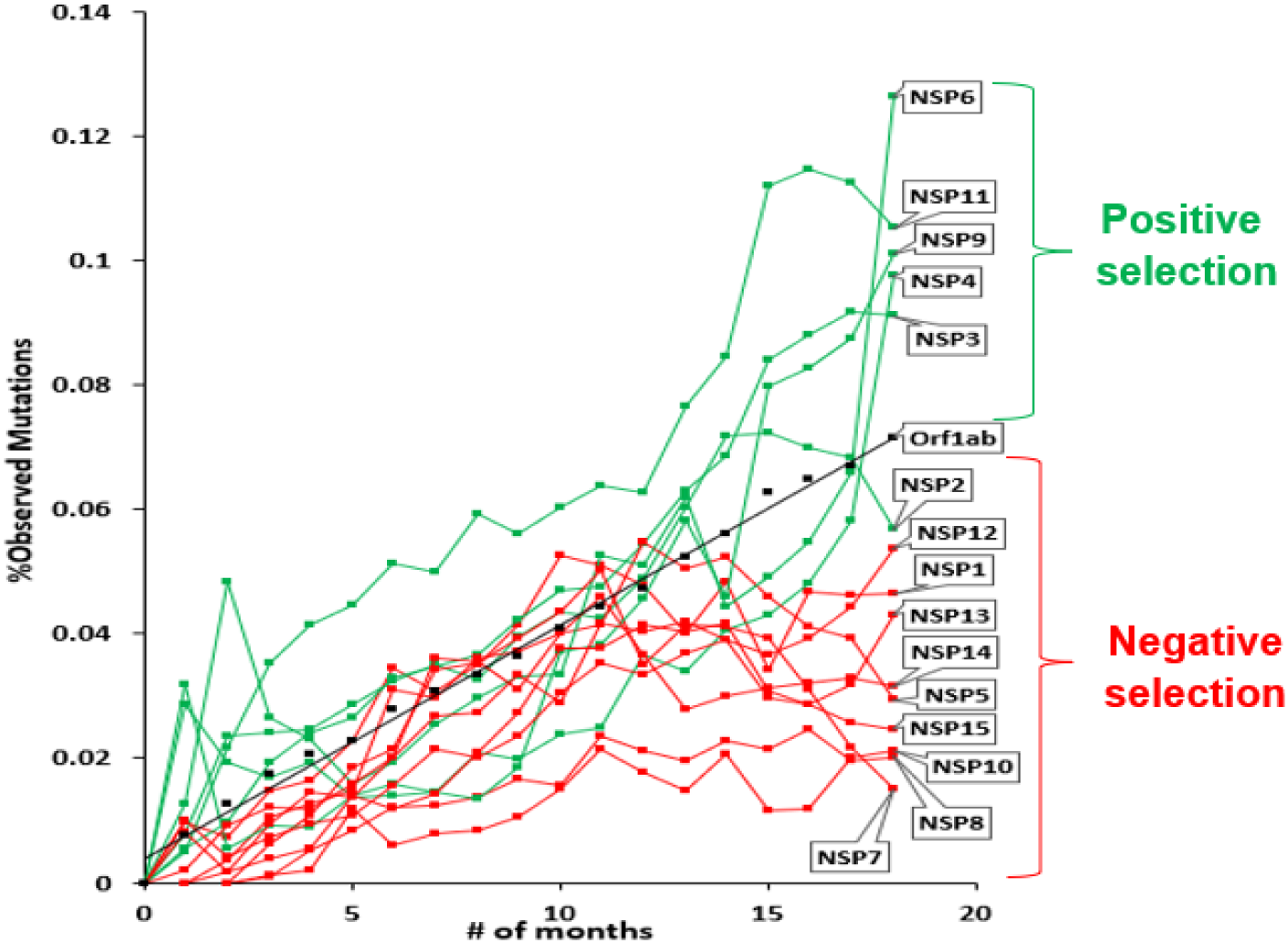
Demonstration of NNBST at the subgenetic level of Orf1ab and NSP1-15. Monthly percent total nucleotide substitution rate for Orf1ab and NSP1-15 genes over 19 months. NSP segments exhibiting faster or slower substitution rates relative to Orf1ab are under relative positive and negative selection with respect to Orf1ab (neutral selection).

Due to the observed molecular clock feature of the Orf1ab SSR (c/µ = 0.11 and R^2^ = 0.9854), the faster and slower SSRs of NSP1-15 due to more positive and negative selection can balance out and produce the Orf1ab SSR, similarly as the individual SSRs produce the GSR. From this, NSP (2, 3, 6, 9 and 11) seem to undergo more positive selection (c/µ > 0.11), whereas NSP (1, 4, 5, 7, 8, 10, 12, 13, 14 and 15) seem to undergo negative selection (c/µ < 0.11). Hence, NNBST can explain the seemingly time-independent total NT substitution rate of Orf1ab.

## Discussion

The c/µ test provides a powerful tool to resolve the “neutralist-selectionist” debate. If the assumption of a time-independent and site-independent mutation rate (µ) is true (**Figure 3**), its time-dependence (molecular clock or not) and site-dependence (SRSRS or PDRSR of c/µ) for a species will directly tell which molecular evolution theory (ST, ONNT, KNT, NNBST and NNUST) does the species follow (**Table 2**).

**Table 2a.**
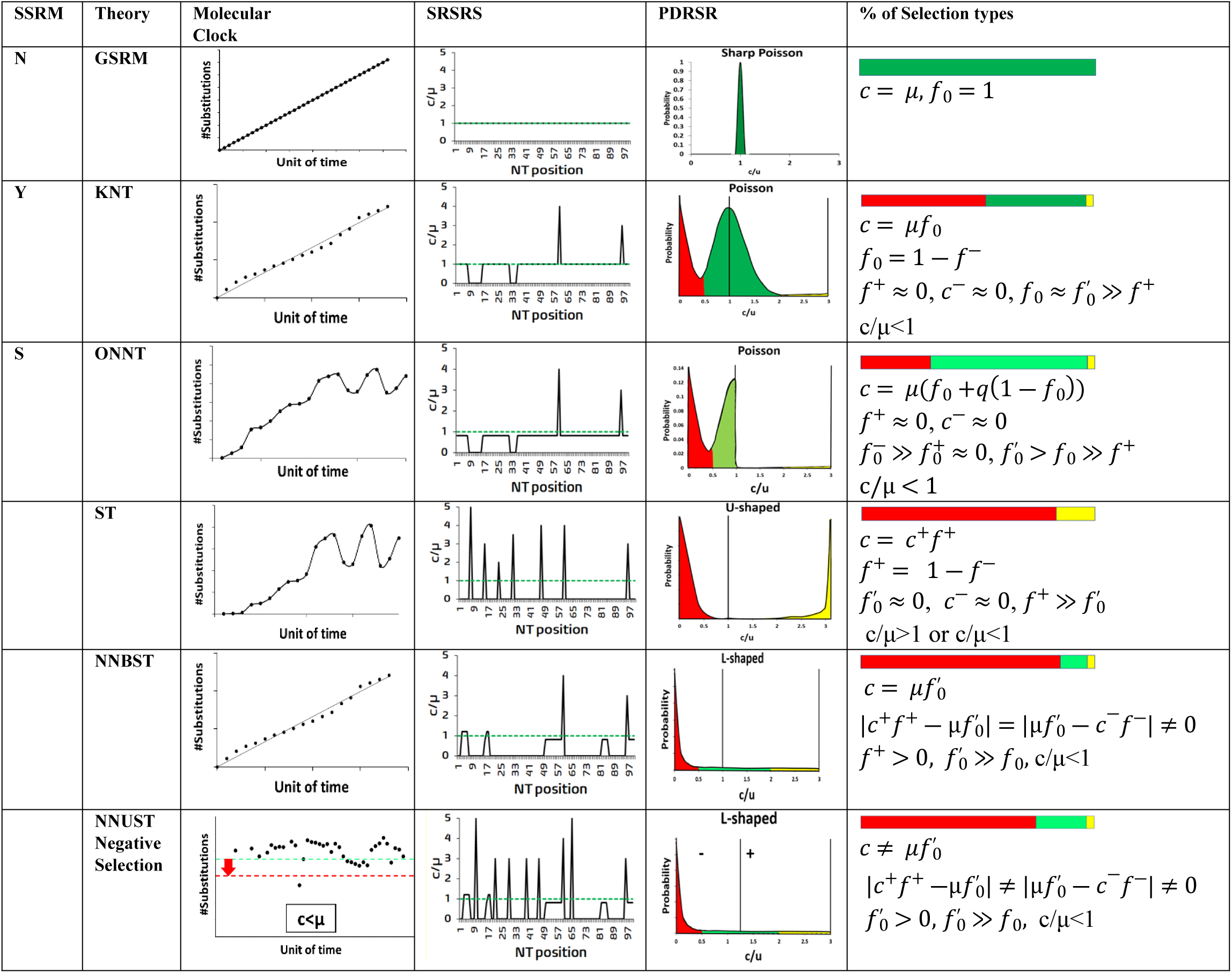
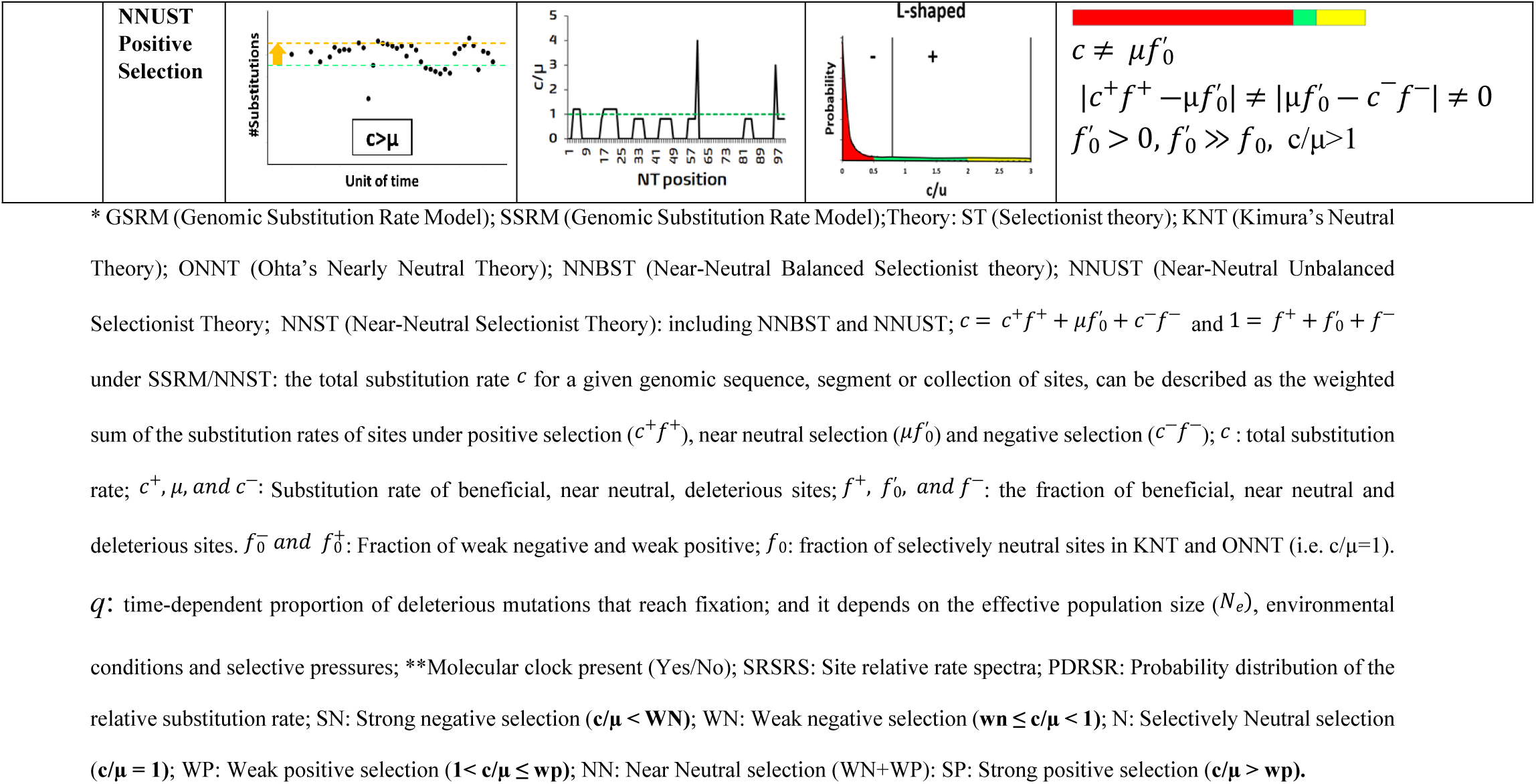
Summary of two substitution rate models (GSRM and SSRM) and five theories (ST, KNT, ONNT, NNBST, NNUST) of molecular evolution in terms of molecular clock, SRSRS, PDRSR and percentage of sites under different selection types (strong negative/red, neutral/green, nearly neutral including (weak negative and weak positive)/light green, and strong positive/dark yellow).

As for SARS-CoV-2, molecular clock was observed only for 24 out of 49 segments (**Figure S2-S3**), and an L-shaped PDRSR was observed for all 49 segments (c/µ_i_) (**Figure S4-S5**), **First,** the dramatic change from the spike-shaped PDRSR of c/µ centered at 1 (*f*_0_ = 1) under GSRM (**Figure 9A**) to the L-shaped PDRSR of c/µ (**Figure 9B)** show the power of natural selection rather than genetic drifting.

**Figure 10.**
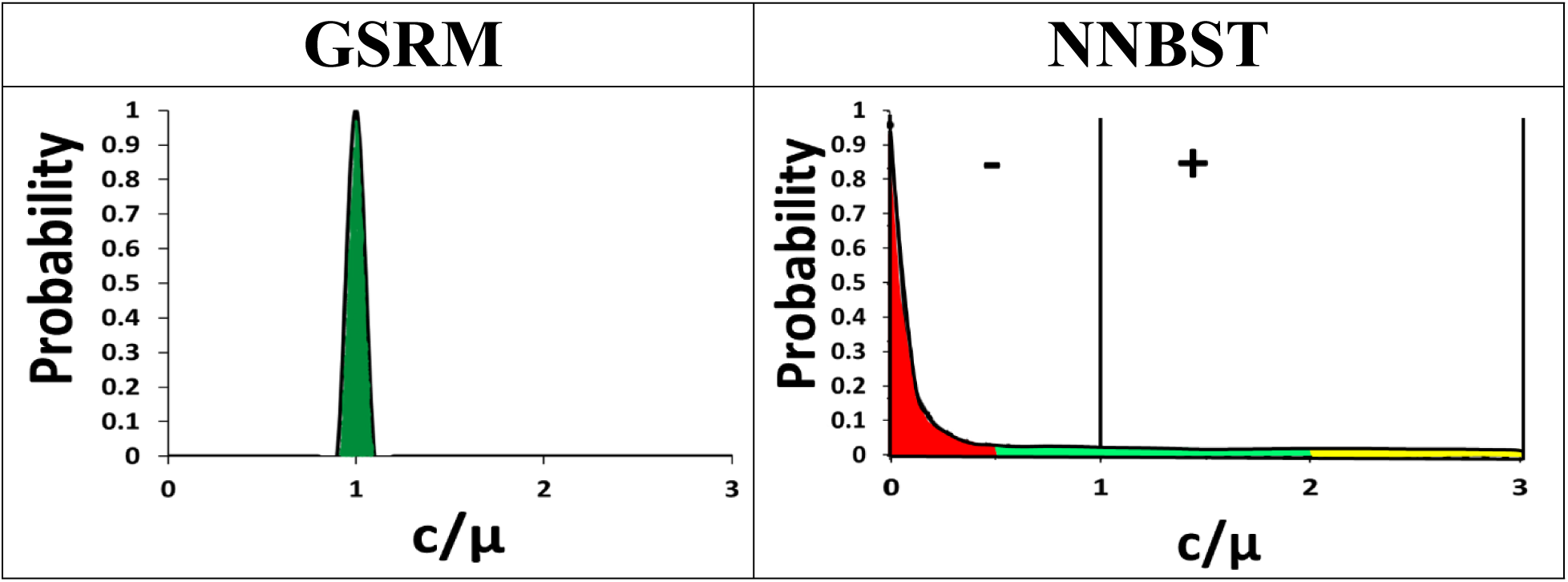
Comparison of the Poisson-shaped PDR of c/µ predicted by GSRM and the L-shaped PDR of c/µ obtained from the SARS-COV-2 genomic datasets explained by NNBST. Selection type is denoted by color (red: negative selection; green: neutral selection; yellow: positive selection).

Indeed, *f*_0_ of this virus’s genome is just 0.03% rather than 100% under GSRM. KNT and ONNT appear to overestimate the fraction of strict neutral mutations and thus the role of genetic drifting. This overestimate is clearly demonstrated by the high ratio ∼577 of near neutral to strictly neutral sites of this virus’s genome. Both natural selection and genetic drifting play important roles in the fixation of near neutral mutations. **Second**, although weak negative sites are typically more abundant than weak positive sites, weak positives do exist. WP:WN ratio of this virus’s genome is ∼5. The early version of ONNT with a shift model predicts only weak negative sites, the later version with a fixed model predicts both weak negative and weak positive sites[16]. Using the simplest form of the fixed model, Gillespie has shown that 50% of the substitutions that become fixed are deleterious and the other 50% are advantageous[17]. **Third**, although weak negative sites are typically more abundant than the positive sites for most of the segments, the exception for a few segments exists. While WN:P ratio of this virus’s genome is ∼9, that of Orflab (0.73) and NSP7 (0.33) is less than 1. While ST might underestimate the important role of genetic drifting in fixing slightly deleterious mutations, KNT and ONNT might underestimate the critical role of positive selection in a few segments and the negative selection in fixing slightly deleterious mutations. **Fourth,** the balance condition (**Equation 8a**) explains molecular clock. This molecular clock is due to nearly neutral mutations rather than strictly neutral mutations under the balance condition. The early version of ONNT predicts a population-size dependent substitution rate, since the lower purification efficiency on slightly deleterious mutations in a smaller population would lead to their higher substitution rate[16]. Yet, as the population size of SARS-COV-2 changes over seven magnitude (10^0^-10^7^), its GSR is still close to constant[13]. If the lower fixation efficiency on slightly beneficious mutations in the small population would lead to a lower substitution rate, then the higher rate from the slightly deleterious mutations will be cancelled out, and thus the population-size independent molecular clock can be observed. This mechanism is allowed in NNBST. **Fifth**, if the balance condition is not satisfied, then molecular clock will not be observed as predicted by NNUST. However, there is no good reason to support the positive selection must be balanced with the negative selection, thus a balanced selection should be less frequent than an unbalanced selection. In fact, lack of a constant molecular clock was observed in 25 of 32 RNA viruses [18]. Our unpublished data show that only ∼5% of 2000 viruses show molecular clock. Put together, we proposed Near-Neutral Selectionist Theory (NNST) to combine ST with ONNT to acknowledge the important roles of both genetic drifting and natural selection in the molecular evolution. As to SARS-COV-2’s genome with a scaled-up µ (5.45-fold of the original one[13]), 84.21%, 14.28%, 1.33% and 0.19% mutations are under strong negative, weak negative, weak positive and strong positive selection, respectively. Of the substitutions that fix in the population, 90.41% (14.28%), 8.41% (1.33%) and 1.19% (0.19%) are slightly deleterious, slightly advantageous and strong advantageous. While genetic drifting and the purifying selection of natural selection plays an important role in fixing 98.82% of near neutral mutations in the population, the positive selection of natural selection also plays an important role in fixing 9.6% of advantageous mutations.

Limitations of c/µ test and future directions are briefly discussed as follows. **1)**. Our assumption of time-independent site-independent global genomic mutation rate (µ) remains to be validated from the replication experiments. **2).** When ancestral sequences are not known, a phylogenetic tree must be inferred from a set of sequences and ancestral sequences with a root sequence must be reconstructed to get the reference for mutation counting. **3).** c must be sampled from a representative population of sufficient size; therefore, it might not capture the founder effect and genetic bottleneck conditions when population size is small. Accurate establishment of c should be obtained from a genomic dataset given **a)** sufficient sample size is obtained, **b)** sufficient evolution time has passed, and **c)** samples are representative of the effective population. Sufficient sample sizes increase the likelihood of representing the effective population by mapping the genetic variation accumulated over evolution time. Moreover, ensuring enough evolution time has passed allows mutations to become fixed in the effective population and the establishment of a molecular clock feature (if possible). Finally, a large, representative sample size can prevent the pitfall of the Founder Effect, which unique mutations and phenotypes associated with breakaway subpopulations may not represent the effective population. **4).** c/µ is the overall indirect fitness measurement for nucleic acid and protein sequence from experimental measurements, but it does not tell the mechanism of fitness change (e.g. selection coefficients) that can be used for predicting the molecular adaption.[14] While the c/µ values for a given segment and NT/AA site are useful for measuring the relative fitness change, additional analysis is required to probe the nature of such change for getting selection coefficients.[14] For instance, molecular modeling (e.g., molecular docking and dynamics simulations) can provide insight into mutation effects on protein/nucleic acid structure and drug binding interactions at the atomic level, and/or be experimentally characterized.

## Conclusion

In this comprehensive analysis of the relative substitution rates (c/µ) across various segments of the SARS-COV-2 genome, we have demonstrated notable departures from traditional evolutionary models. By extending our investigation to 26 translated regions, 12 untranslated regions, and 10 transcriptional regulatory sequences, we observed L-shaped probability distributions of c/µ and variable presence of molecular clocks, supporting both our Near-Neutral Balanced Selectionist Theory (NNBST) and Near-Neutral Unbalanced Selectionist Theory (NNUST). Our findings significantly challenge the existing paradigms of evolutionary biology by demonstrating that neither strictly neutral mutations nor solely beneficial mutations dominate the genomic landscape. Instead, nearly neutral mutations including both slightly deleterious and slightly beneficious, influenced by both genetic drift and natural selection, predominate. This nuanced understanding suggests that the fixation of mutations is influenced by a balance of selective pressures rather than by simple stochastic events or direct selection alone. The discovery of L-shaped distributions across all studied segments, coupled with the variable presence of molecular clocks, underscores the dynamic nature of viral evolution and the need for a revised theoretical framework. The NNST (Near-Neutral Selectionist Theory) that we propose offers such a framework, combining elements of traditional neutral and selectionist theories to more accurately reflect the evolutionary processes observed in SARS-COV-2 and potentially other RNA viruses. Our research points to the necessity for a reevaluation of the “neutralist-selectionist” debate, advocating for a more integrated approach that considers the spectrum of mutation effects from deleterious to beneficial. This approach not only enhances our understanding of viral evolution but also may inform future therapeutic strategies by highlighting the evolutionary potential and constraints within viral genomes. As we continue to refine our theoretical models and expand our empirical datasets, it becomes increasingly clear that the molecular mechanisms governing evolution are far more complex and interconnected than previously assumed. The work presented here marks a significant step towards resolving long-standing debates in evolutionary biology and sets the stage for future investigations that will further elucidate the intricate dance of mutation, selection, and drift in shaping the genomes of living organisms.

## Methods

A replication-selection model for describing molecular evolution of a virus, definition of the substitution-mutation rate ratio (c/µ) to quantify the fitness change due to mutations, use of c/µ probability distribution to define different evolution empirical models including Genomic Substitution Rate Model (GSRM), Segment Substitution Rate Model (SSRM); and evolutionary theories including ST, KNT, ONNT and NNBST, and the pipelines to calculate c/µ and its change over time, have been described in our previous paper[13]. In this paper, we concisely describe them (**Table 1 and 2);** and focus on deriving the general equations to define them.

**Table 1.**
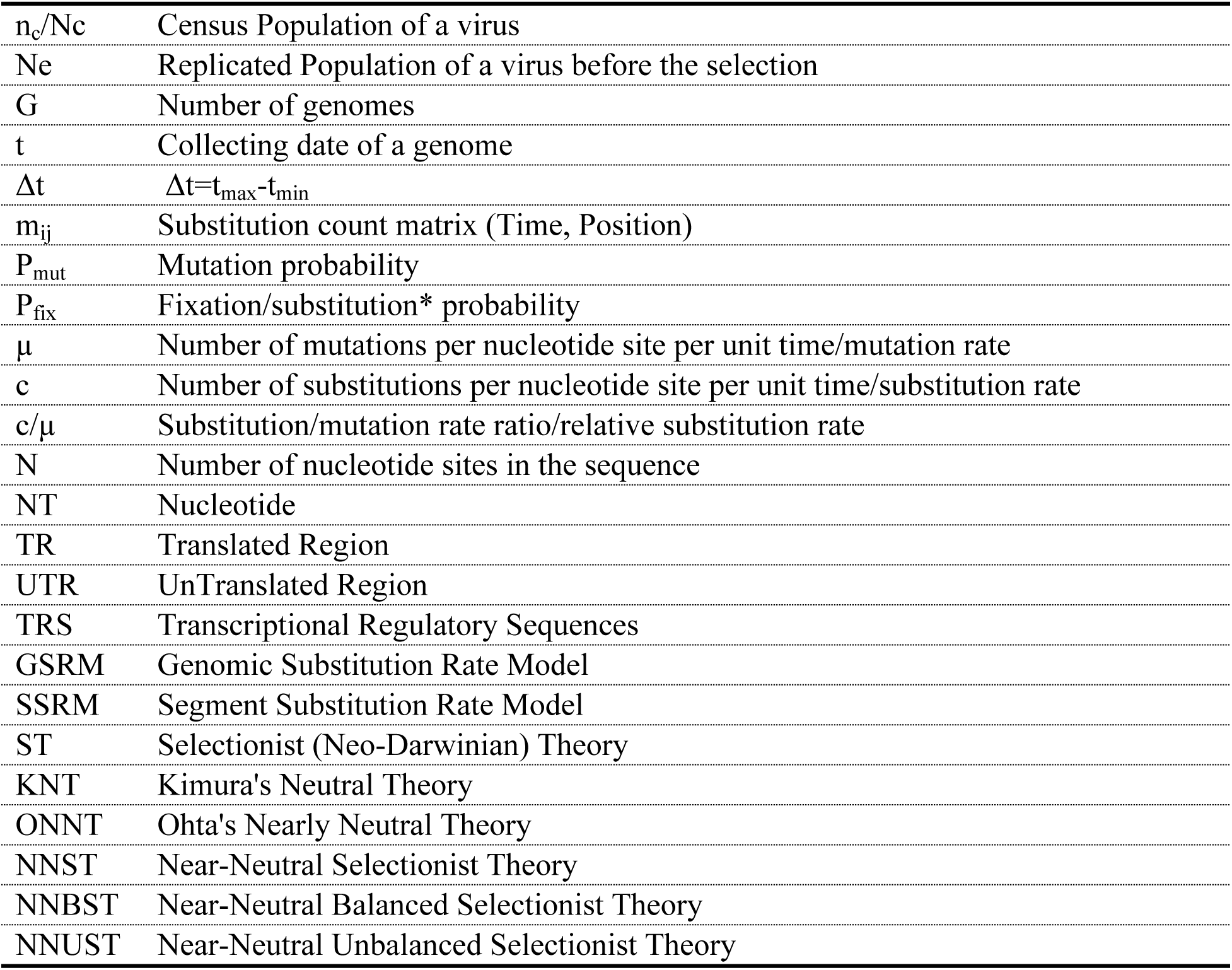
Definitions of Major symbols.

|  |  |
| --- | --- |
| $n_c/N_c$ | Census Population of a virus |
| $N_e$ | Replicated Population of a virus before the selection |
| $G$ | Number of genomes |
| $t$ | Collecting date of a genome |
| $\Delta t$ | $\Delta t = t_{\max} - t_{\min}$ |
| $m_{ij}$ | Substitution count matrix (Time, Position) |
| $P_{\text{mut}}$ | Mutation probability |
| $P_{\text{fix}}$ | Fixation/substitution* probability |
| $\mu$ | Number of mutations per nucleotide site per unit time/mutation rate |
| $c$ | Number of substitutions per nucleotide site per unit time/substitution rate |
| $c/\mu$ | Substitution/mutation rate ratio/relative substitution rate |
| $N$ | Number of nucleotide sites in the sequence |
| NT | Nucleotide |
| TR | Translated Region |
| UTR | UnTranslated Region |
| TRS | Transcriptional Regulatory Sequences |
| GSRM | Genomic Substitution Rate Model |
| SSRM | Segment Substitution Rate Model |
| ST | Selectionist (Neo-Darwinian) Theory |
| KNT | Kimura's Neutral Theory |
| ONNT | Ohta's Nearly Neutral Theory |
| NNST | Near-Neutral Selectionist Theory |
| NNBST | Near-Neutral Balanced Selectionist Theory |
| NNUST | Near-Neutral Unbalanced Selectionist Theory |

### The replication-selection model to describe molecular evolution under natural selection, the relative substitution rate (c/µ) to quantify the fitness change of mutations, the discrete c/µ probability distribution to define different models/theories of molecular evolution

A replication-selection model was introduced to describe the evolution of a virus in **Figure 2** of our previous paper[13]. An elaborated version has been generated (**Figure 2A**) and will be described here. The numbers used to represent different viral populations are arbitrary and are kept small here for simple calculations; more realistic conditions likely contain viral populations several orders of magnitude higher. A nascent virus census population containing n members (n_c_=3) undergoes viral replication with a mutation rate (µ) that induces mutations at three different sites with equal probability (P_mut_ = 3/12, 3/12, 3/12) over the time period (t). After replication, a mutant population is produced (N_e_ = 12) and is subjected to either positive selection (yellow box), neutral selection (green box) or negative selection (red box) on the first NT position from the host (e.g., host immune response, host dies, vaccinations). The new population surviving selection is observed (N_c_ = 4) and bears fixed mutations with a probability for the three different mutants under positive selection (P_fix_ = 2/4, 1/4, 1/4), neutral selection (P_fix_ = 1/4, 1/4, 1/4) and negative selection (P_fix_ = 0/4, 1/4, 1/4) for the first NT position. The process is cyclic and continues for successive viral generations within the host and following infections with other hosts. It must be noted that the population N_c_ is used for genomic and mutation analyses, whereas population N_e_ is much harder to observe unless an in vitro experiment is conducted. Although if performed, µ could be experimentally determined and provide a more accurate c/µ per position and genomic segment.

One nuance to this process is that the total population count before and after selection can change, but the relative abundance of different genotypes should stay the same under neutral selection (i.e. an equal fixation probability (1/N_e_) for each individual virus in the replicated population N_e_).[4] For example, a hypothetical population after mutation (N_e_ = 12) contains 3 wild-type sequences (3/12 or 1/4), 3 red mutants, 3 green mutants and 3 blue mutants, corresponding to their relative abundance (1:1:1:1). After neutral selection, only 25% of the sequences survive (N_c_ = 4) of 1 wild-type, 1 red mutant, 1 green mutant and 1 blue mutant (**Fig. 2A** green box), however the relative abundance is the same (1:1:1:1). Therefore, P_mut_ = P_fix_ = ¼ for red mutation, thus, µ = c by dividing the same time period (t). This relationship is general so that when the rate of mutation generation (µ) is equal to the rate of substitution (c) in the population suggests neutral selection without a preference (c/µ=1).[4] Only when neutral selection is well defined, then natural selection can be defined. If the substitution rate of the red mutation is higher than the rate of its generation (c/µ>1), the mutation must be beneficial to increase the fitness of the virus (positive selection **Fig2A**, yellow box). If the substitution rate of red mutation is lower than the rate of its generation (c/µ<1), the mutation must be deleterious (i.e. negative selection **Fig. 2A**, red box). Therefore, c/µ can quantify the fitness change of mutations at a specific NT site of a genome.

To define µ, two prior assumptions are made in the replication step of our replication-selection model (**Figure 3**).

**Figure 3.**
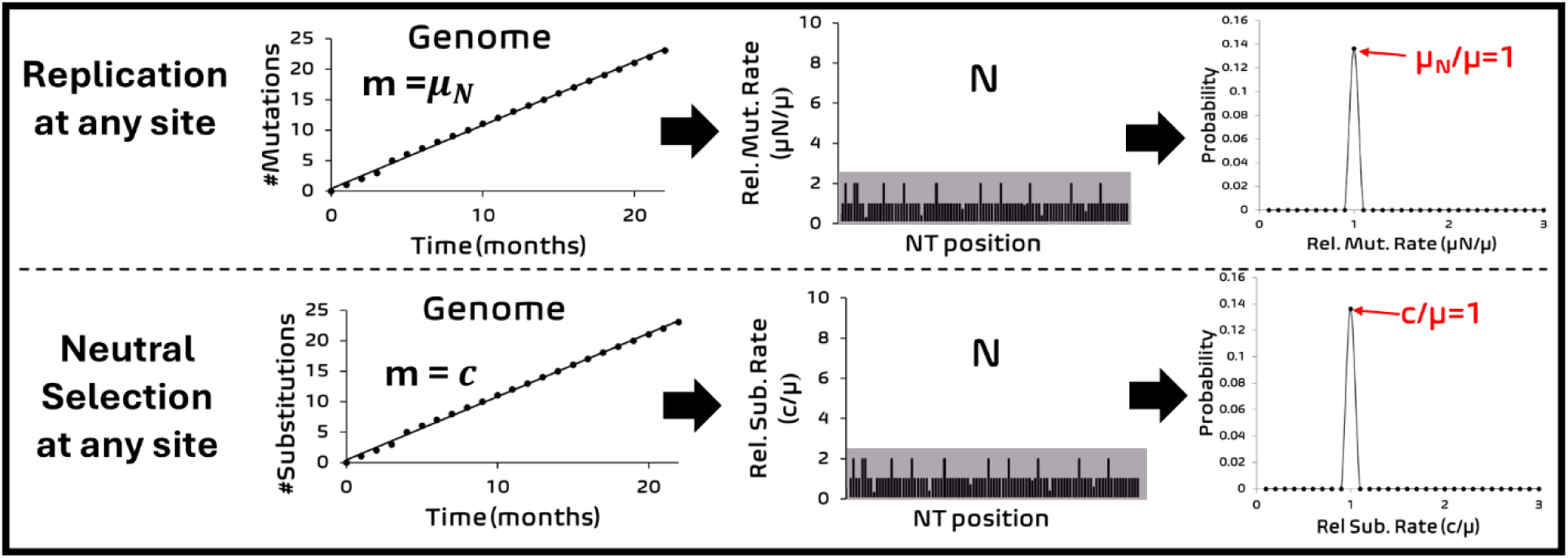
The viral genomic sequence under neutral selection. During replication, the viral polymerase induces copy errors at rate µ at each nucleotide site in the copied viral sequence strand. The total nucleotide sequence length (N) is the sum of all nucleotide sites in the translated and untranslated regions. Mutations are fixed into each site at rate c. The resulting probability distribution of the relative mutation rate (μ_N_/μ) for a given site under neutral selection is described as a Poisson distribution curve.

For viral replication, 1) the viral replication machinery (e.g., polymerase with/without proofreading enzymes) has no preference for where it induces copy error and does so at any NT site with a phenomenological rate μ. 2) The polymerase structure does not significantly change from induced mutations; thus, the polymerase error rate (μ) does not significantly change over the measuring time period (i.e. time-independent constant leading to strict molecular clock in **Figure 3 left top panel**). Nonetheless, these two prior assumptions are shared among many mutation models[14, 19].

Key features of the discrete c/µ probability distribution define different models/theories of molecular evolution (**Table 2**). The c/µ of each NT site in a genome of c/µ (defined as Site-Relative Substitution Rate Spectrum (SRSRS)) spells out the selection pressure of the mutations at the NT site. SRSRS (**Figure 3 middle**) can be equivalently transformed into a discrete Probability Distribution of Relative Substitution Rate (PDRSR) for all NT sites of the genome including sampling errors (**Figure 3 right**). Assuming all NT sites undergo strictly neutral selection only (i.e. the fraction of strictly neutral sites is one, *f*_0_ = 1; Genomic Substitution Rate Model/GSRM[13]; the Strict neutrality hypothesis [11]), the observed mutation rate at any NT site *j* (substitution rate, c_j_) is equal to the spontaneous mutation rate in individual organisms (*c* = μ **Figure 3**) [3, 4]. Therefore, the substitution rate for any genetic segment, UTR, TRS, sub-genetic segment is a constant that is equal to the spontaneous mutation rate (μ), leading to a global molecular clock.[2] The temporal and site features outputted from the mutation model in the mutation step will be observed again under neutral selection **(Figure 3 Bottom Panel**). Put together, empirical hallmarks of GSRM are the global molecular clock, discrete almost uniform distribution of c/µ at any site of the genome, and Poisson distribution centered at c/µ=1 with a broadened peak due to random sampling error (**Table 2**).

The Segment Substitution Rate Model was introduced to allow the time-dependent natural selection pressure acting on the genome, thus causing the deviation of the substitution rate of a specific segment or a NT site *j* from the constant mutation rate (i.e. c_j_/µ≠1). Various theories have been proposed to capture the relative contribution between genetic drifting forces (neutral selection) and natural selection forces in determining the evolutionary substitution rate. In KNT (**Table 2**), while most NT sites are under negative selection (*f*^―^ > *f*_0_; *f*^―^: the fraction of deleterious sites), the sites under strictly neutral selection (c/µ=1) are much more abundant than the sites under positive selection (c/µ>1, *f*^+^ ≈ 0, *f*_0_ ≫ *f*^+^; *f*^+^: the fraction of beneficious sites) [20]. The hallmark of KNT is the molecular clock due to large number of neutral sites and a Poisson distribution of centered at c/µ=1 (**Table 2**). Along the same line of KNT, ONNT [7, 8] predicts while most NT sites are under negative selection (*f*^―^ > *f*_0_), the sites under strictly neutral selection (c/µ=1) or nearly neutral selection (c/µ∼1, *f*^′^_0_:the fraction of near neutral sites) aremuch more abundant than the sites under positive selection (*f*^+^ ≈ 0, *f*^′^_0_ > *f*_0_ ≫ *f*^+^). Hallmarks of ONNT are population-size dependent substitution rate (**Eqn 6**: *c* = μ(*f*_0_ +*q*(1 ― *f*_0_); *q*: time-dependent proportion of deleterious mutations that reach fixation, which is dependent on the effective population size (*N*_e_), environmental conditions and selective pressures[11]) and an asymmetric distribution around c/µ=1 due to mostly slightly deleterious near neutral mutations (**Table 2**). In ST [2], NT positions largely exhibit natural selection (c_j_/µ≠1) and few sites under strictly neutral selection (**Table 2**). Hallmarks of ST are time-dependent substitution rate, and a bimodal distribution of c/µ showing one peak of negative selection (c/µ<1) and the other peak of positive selection (c/µ>1). In NNBST, hallmarks are L-shaped distribution of c/µ and molecular clock due to the cancelling out of the rate of change of the sites under more positive selection by that of the sites under more negative selection (defined as a balance condition that will be derived later). The balance condition is more general than the original assumptions in KNT that beneficial sites are so rare. In NNUST, hallmarks are L-shaped distribution of c/µ and lack of molecular clock due to the failure of the balance condition (**Table 2**). Again, NNUST is more general than ST without assuming that nearly neutral sites are so rare. If the average substitution rate is faster than the mutation rate, positive selection dominates the overall selection. Otherwise, the average substitution rate is slower than the mutation rate, thus negative selection dominates the overall selection.

### While the balance condition leading to molecular clock in NNBST allows larger fraction of positive mutations than KNT does, the failure of the balance condition leading to lack of molecular clock allows a larger fraction of nearly neutral mutations than ST does

The contribution of each selection/mutation type towards the total substitution rate for a given genomic sequence, segment or collection of sites, can be described as the weighted sum of the substitution rates of sites under strong positive selection (*c*^+^*f*^+^), near neutral selection (μ*f*^′^_0_) and strong negative selection (*c*^―^*f*^―^) following NNST:

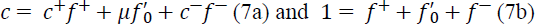

Although the second neutral site term could be time-independent where the mutation rate (μ) is constant and *f*^′^_0_ could be time-independent constant, the overall substitution rate is more likely to be time-dependent because of time-dependence of the first strong positive site term and the third strong negative site term. However, the total rate can be time-independent under the balance condition of NNBST where the increase of the rate due to the advantageous mutations (*c*^+^*f*^+^, *c*^+^ > µ) is exactly canceled by the decrease of the rate due to the deleterious mutations (*c*^―^*f*^―^, *c*^―^ < µ):

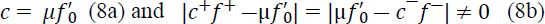

Thus, *f*^′^_0_ (*f*^′^_0_ ≤ 1) and thus *c* is time-independent rate constant.

However, when the balance condition is not satisfied,

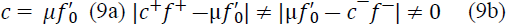

**Equations 9a and 9b** define NNUST, predicting a time-dependent *f*^′^_0_ and *c*. In addition, the substitution rate *c* that can be higher than the mutation rate under effective positive selection (*f*^′^_0_= c/μ>1) or under effective negative selection (*f*^′^_0_ = c/μ<1).

NNBST can be reduced to KNT, which defines the overall substitution rate as the product of the mutation rate with the fraction of strictly neutral mutations only by assuming the fraction of beneficial mutations and the substitution rate of deleterious mutations are negligible(*f*^+^ ≈ 0 and *c*^―^ ≈ 0) and strictly neutral sites comprises the majority of the near neutral mutations (*f*^′^_0_ ≈ *f*_0_):[6]

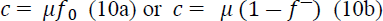

Conversely, NNUBT can be reduced to ST, which define the overall substitution would be the product of the substitution rate of strong positive sites (*c*^+^) with its fraction (*f*^+^) by assuming the fraction of neutral mutations and the substitution rate of deleterious mutations are negligible small (*f*^′^_0_ ≈ 0 and *c*^―^ ≈ 0):

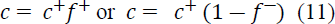

In other words, only beneficial mutations can be fixed in the population.

First, the balance condition in NNBST allows a non-zero fraction of positive selection sites than KNT so that molecular clock is not sufficient to prove *f*_0_ > *f*^+^. NNUST allows a more non-zero fraction of neutral selection sites than ST so that lack of molecular clock is not sufficient to prove *f*^+^ > *f*_0_.

Second, while **equation 8a** of NNBST or equation **10a** of KNT predicts that a molecular clock with variable *f*^′^_0_ or *f*_0_ of different genome segments under negative selection only and the maximum substitution rate is limited to the mutation rate (*c*→*u* when *f*^′^_0_ or *f*_0_→1) due to lack of the positive selection; **equation 9a** of NNUST and **11** of ST predicts a time-dependent substitution rate due to time-dependent selections: (*c*^+^*f*^+^+*c*^―^*f*^―^) in NNUST and (*c*^+^*f*^+^) in ST, leading to either an effective negative selection with a upper rate limit of *u* or an effective positive selection without a upper rate limit.

### Position-based and time-based approaches for calculating substitution rates (c)

Given the reference genome and a set of genomes for a species, the number of substitutions at each NT site can be counted for a given time interval after multiple sequence alignment using MAFFT.[21] To calculate a substitution rate (substitutions per NT per unit time, c) from a reference sequence of length N and an aligned sequence set of size G, a two-dimension (Time x Position) substitution count matrix (*m_ij_*) at various collection dates and at different genome positions was obtained, but there are two ways for averaging substitution frequency across position and time: time-based and position-based approaches (**eq. 12**). While the time-based approach (eq. 12 left) begins by averaging substitution counts across positions to get temporal (timeline) trends, and then across time (Δ*t* = *t_max_* ― *t_min_*), the position-based approach (eq. 10 right) averages substitution counts across time to get SRS, and then across positions.

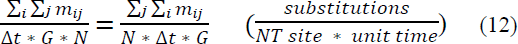

Both methods offer distinct perspectives on substitution dynamics: while a timeline from the time-based approach emphasizes temporal trends (**Fig1 E)**, SRSRS form the position-approach highlights spatial patterns (**Fig1 C**). Both approaches should give identical results of an absolute rate, but we modified the time-based approach to allow a least square fitting (y=a*x, R^2^) to timeline for getting the final rate. This rate should not be identical to the rates calculated from the position-based approach. This fitting implicitly assumes molecular clock of a constant rate (**Equations 1-5)**. If this assumption is true (*m_i_* = *rate* ∗ *t_i_*), then the fitting could improve the accuracy of the rates by removing random “noise” in the sampling; and the obtained absolute true rate value differs from that from the position-based approach by a constant 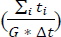. The scaling constant will be canceled when calculating the relative rates (c/µ), therefore both ways should give very similar values if the molecular clock assumption is true. The coefficient of determination (R^2^) from the fitting can offer a statistical measure on how good the molecular clock assumption holds. If R^2^ is too small (R^2^ ≤ 0.6000), the rates from the position-based approach should be used. Nonetheless, SRSRS is useful to identify adaptive mutations. SRSRS can be converted into PDRSR for each gene/UTR and the whole genome for characterization of different evolution theories (**Table 2**).

### The proportion of the strictly neutral, nearly neutral, beneficial, deleterious mutations can be determined from PDRSR. The fraction of strictly neutral mutations 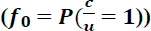 can be precisely obtained from PDRSR. The fraction of near neutral mutations was determined through equation 8a/9a 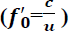 for the segments with molecular clock and for the segments without molecular clock under an effective negative selection (c/μ<1), leading to determination of its weak negative boundary (WNB), its weak positive boundary (WPB=1/WNB); the fraction of mutations under strong negative (SN), weak negative (WN), weak positive (WP) and strong positive (SP) selection; and ratio of mutations under weak negative to positive selections (WN:P) and ratio of near neutral to strictly neutral mutations (*f*′_0_:*f*_0_).

In our previous paper [13]**, WNB is arbitrarily set to ½ and so is WPB (2=1/WNB). Given equation 8a/**9a (*c* = μ*f*′_0_), WNB can be determined from the fraction of nearly neutral mutations (*f*’_0_) by finding 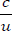 such that its cumulative probability is equal to 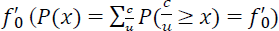 in this study. WPB is arbitrarily set to 1/WNB. Given the cumulative probability at conserved sites 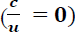, WNB, strictly neutral 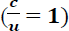, and WPB, the fraction of mutations under strong negative 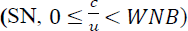 weak negative 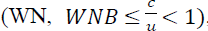, weak positive, 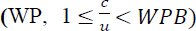 strong positive selection 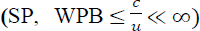 can be obtained from the cumulative probability function. While the fraction of near neutral (*f*′_0_) mutation is equal to the sum of mutation under WN and WP selection; the fraction of positive (P) mutation is the sum of the mutations under WP and SP selection. The ratio of the mutations under weak negative to positive selection (WN:P) and the ratio of near neutral to strictly neutral mutations (*f*′_0_:*f*_0_) were calculated to validate the theories. *f*′_0_, R2, WNB, N, WPB, SN, WN, *f*_0_, WP, SP, WN:P and *f*′_0_:*f*_0_ for the 24/25 segments with/without molecular clock are listed in Table 3 and 4, respectively.

## Acknowledgements

C.W acknowledges the support by the New Jersey Health Foundation (PC 76-24) and the National Science Foundation under Grants NSF ACI-1429467/RUI-1904797, and ACESS/BIO230145. The Anton2 machine at the Pittsburgh Supercomputing Center (PSCA170090P) was generously made available by D. E. Shaw Research.

## Author Contributions

Conceptualization and Equation Derivation, C.W.; MatLab scripts, C.W. and N.J.P.; Data preprocessing, N.J.P.; Plots and Tables, N.J.P. and C.W.; writing—original draft preparation, review and editing, C.W. and N.J.P.; funding acquisition, C.W.. All authors have read and agreed to the published version of the manuscript.

---

Table S1. c total substitution rate slope and c R^2^ values for each dataset and averaged over each dataset for all SARS-COV-2 UTR and TRS segments.

Table S2. Time-based c total genomic substitution rate slope and c R^2^ values for each dataset and averaged over each dataset for all SARS-COV-2 coding proteins.

Table S3. Position-based c/µ (c/µ^a^), time-based c/µ (c/µ^b^) and their absolute difference values for the All-UTR, All-TRS and each UTR and TRS over 19 months, combined from Datasets A1A-A1C.

Table S4. Time-based c/µ (c/µ^a^), Position-based c/µ (c/µ^b^), their absolute and percent differences for the genome, All-UTR, All-TR and each coding segment of SARS-COV-2 over 19 months, combined from Datasets A1A-A1C.

Table S5. Abundance of sites under different selection types for segments exhibiting molecular clock feature. (Column 1): Relative abundance of sites under WN, WP and SP selection. (Column 2): Abundance of sites under NN and P selection. (Column 3): Abundance of sites under SN, NN and SP selection. (Column 4): Abundance of sites under WN and SP selection. (Column 5): Abundance of sites under N and P selection. See **Figures 7 and S6** for graphical representations.

Table S6. Abundance of sites under different selection types for segments exhibiting non-molecular clock feature. (Column 1): Relative abundance of sites under WN, WP and SP selection. (Column 2): Abundance of sites under NN and P selection. (Column 3): Abundance of sites under SN, NN and SP selection. (Column 4): Abundance of sites under WN and SP selection. (Column 5): Abundance of sites under N and P selection. See **Figures 7 and S6** for graphical representations.

Table S7. Tabulated parameters for the WN:P to c/μ relationship in **Figure 8**.

Table S8. Tabulated parameters for the fo’:fo to c/µ scaling factor relationship in **Figure 8**.

Figure S1. Percent genomic variation of sequences in set A1a with the references sequences Wuhan-Hu-1 (left) and Wuhan IPBCAMS-WH-01 2019 (right) set.

Figure S2. The percent total NT substitution rate for each TR, UTR and TRS segment over evolution time and averaged over the three combined datasets. See **Table S1-S2** for tabulated regression parameters.

Figure S3. The total percent NT variation for the genome, All-TR, All-UTR, All-TRS and each coding and non-coding gene segment over evolution time for each individual dataset. See **Table S1-S2** for tabulated regression parameters.

Figure S4. c/µ Empirical probability distribution function of 24 segments exhibiting molecular clock feature.

Figure S5. c/µ Empirical probability distribution function of 25 segments exhibiting non molecular clock feature.

Figure S6. c/µ Cumulative probability distribution function 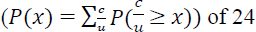 segments exhibiting molecular clock feature, showing the abundance of sites under different selection types. The boundaries for weak negative (red line) to neutral selection (green line) to weak positive selection (orange line) and their determined c/µ positions are noted. Strong negative selection and strong positive selection would be to the left and right of the red and orange lines, respectively. A broken lined leading on the x-axis represents the distance between c/µ = 3.0 to the weak positive selection boundary. See Table 3 for tabulated c/µ boundaries and percent abundances for each selection type.

Figure S7. c/µ Cumulative probability distribution function of 25 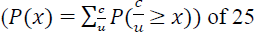 segments not exhibiting molecular clock feature, showing the abundance of sites under different selection types. The boundaries for weak negative (red line) to neutral selection (green line) to weak positive selection (orange line) and their determined c/µ positions are noted. Strong negative selection and strong positive selection would be to the left and right of the red and orange lines, respectively. A broken lined leading on the x-axis represents the distance between c/µ = 3.0 to the weak positive selection boundary. See Table 3 for tabulated c/µ boundaries and percent abundances for each selection type.

Figure S8. Percent selection types of each segment exhibiting molecular clock features decomposed into three categories for strong negative, near-neutral and strong positive selection (top row), weak negative and strong positive selection (middle row) and negative and positive selection (bottom row) using a c/µ scaling of 5.46. See **Table S5** for percent selection type values.

Figure S9. Percent selection types of each segment exhibiting non-molecular clock features decomposed into three categories for strong negative, near-neutral and strong positive selection (top row), weak negative and strong positive selection (middle row) and negative and positive selection (bottom row) using a c/µ scaling of 5.46. See **Table S6** for percent selection type values.

